# EYES ABSENT and TIMELESS integrate photoperiodic and temperature cues to regulate seasonal physiology in *Drosophila*

**DOI:** 10.1101/2020.03.05.979682

**Authors:** Antoine Abrieux, Yongbo Xue, Yao Cai, Kyle M. Lewald, Hoang Nhu Nguyen, Yong Zhang, Joanna C. Chiu

**Affiliations:** Department of Entomology and Nematology, College of Agricultural and Environmental Sciences, University of California, Davis, CA 95616, USA; Department of Biology, University of Nevada Reno, Reno, NV 89557, USA

**Keywords:** photoperiod, temperature, seasonal timing, alternative splicing, reproductive dormancy, diapause, *Drosophila*

## Abstract

Organisms possess photoperiodic timing mechanisms to anticipate variations in day length and temperature as the seasons progress. The nature of the molecular mechanisms interpreting and signaling these environmental changes to elicit downstream neuroendocrine and physiological responses are just starting to emerge. Here, we demonstrate that in *Drosophila melanogaster*, EYES ABSENT (EYA) acts as a seasonal sensor by interpreting photoperiodic and temperature changes to trigger appropriate physiological responses. We observed that tissue-specific genetic manipulation of *eya* expression is sufficient to disrupt the ability of flies to sense seasonal cues, thereby altering the extent of female reproductive dormancy. Specifically we observed that EYA proteins, which peak at night in short photoperiod and accumulate at higher levels in the cold, promote reproductive dormancy in female *D. melanogaster*. Furthermore, we provide evidence indicating that the role of EYA in photoperiodism and temperature sensing is aided by the stabilizing action of the light-sensitive circadian clock protein TIMELESS (TIM). We postulate that increased stability and level of TIM at night under short photoperiod together with the production of cold-induced and light-insensitive TIM isoforms facilitate EYA accumulation in winter conditions. This is supported by our observations that *tim* null mutants exhibit reduced incidence of reproductive dormancy in simulated winter conditions, while flies overexpressing *tim* show an increased incidence of reproductive dormancy even in long photoperiod.

**Significance Statement:** Extracting information on calendar time from seasonal changes in photoperiod and temperature is critical for organisms to maintain circannual cycles in physiology and behavior. Here we found that in flies, EYES ABSENT (EYA) protein act as a seasonal sensor by adjusting its abundance and circadian phase in response to changes in photoperiod and temperature. We show that the manipulation of EYA levels is sufficient to impair the ability of female *Drosophila* to regulate seasonal variation in reproductive dormancy. Finally, our results suggest an important role of the circadian clock protein TIMELESS (TIM) in modulating EYA level through its ability to measure night length, linking the circadian clock to seasonal timing.

## Introduction

As in plants and other animals, insects possess endogenous photoperiodic timers, which prompt them to undergo physiological and behavioral changes to survive through unfavorable periods. Arguably the most well recognized seasonal response in insects is the induction of overwintering diapause. This phenomenon can be induced at different life stages and is characterized by a reversible arrest in growth and/or reproduction in response to decreasing day length. Since photoperiodic time measurement (PPTM) is critical to seasonal adaptation in insects, it has been studied extensively (1, 2). Yet, the molecular and neuronal basis of the insect photoperiodic timer remain poorly understood.

Extensively studied for its role in eye development (3, 4), the EYES ABSENT (EYA) protein represents a promising target to unravel the molecular mechanisms underlying PPTM. This cotranscription factor with phosphatase activity is highly conserved in the animal kingdom (5) and has been implicated in diverse biological processes such as organ development, innate immunity, DNA damage repair, angiogenesis, and cancer metastasis (6–10). Interestingly, the mammalian ortholog, EYES ABSENT 3 (EYA3) was recently implicated in sheep as a clock-regulated transcription factor in the pituitary gland to promote the transcription of *TSHβ*, leading to an increase in thyroid-stimulating hormone (TSH) and an induction of summer phenotypes in long photoperiod (11–15). This elegant work suggests that EYA3 plays a central role in the photoperiodic switch synchronizing circannual rhythms in reproduction with the environment. Nonetheless, due to inherent limitation of using sheep as animal subjects for those studies and the pleiotropic functions of EYA during development, there is still a lack of *in vivo* functional data from genetic manipulation of *eya* expression to confirm this hypothesis. Here we take advantage of the versatile genetic tools available in *Drosophila melanogaster* to investigate the role of EYA in insect photoperiodism.

Unlike many insect species relying mainly on photoperiodic signal to overwinter (16), *D. melanogaster* requires cold temperature to enhance reproductive dormancy under short photoperiod (17). Whether *D. melanogaster* represents a suitable model to study diapause has been under debate. However recent studies suggest that the photoperiodic component of *D. melanogaster* might be more robust than previously described (18, 19). We found that newly emerged females reared at 10°C for 28 days, exhibit significantly smaller ovaries when exposed to short photoperiod (SP 8L:16D) compared to long photoperiod (LP 16L:8D). Using these conditions as readout for reproductive dormancy in combination with the inducible gene-switch driver to genetically manipulate *eya* expression in a spatiotemporal manner (3, 20–22), we provide functional evidence for the role of EYA in insect PPTM, specifically in promoting winter physiology. We are referring to the phenomenon of ovary growth arrest in *D. melanogaster* as reproductive dormancy rather than diapause due to its reliance on temperature in addition to photoperiodic cues.

Finally, we present results suggesting that the function of EYA in seasonal adaptation is aided by cold temperature-dependent induction of light-insensitive TIMELESS (TIM) isoforms, which contribute to EYA stabilization in winter condition. Our results directly link a key circadian clock protein to insect seasonal timing.

## Results

### Reproductive dormancy is a robust phenotypic readout for winter physiology in *Drosophila melanogaster*

To verify that reproductive dormancy (i.e. ovary size) in *D. melanogaster* can serve as a robust phenotypic readout for long photoperiod (LP 16L:8D) versus short photoperiod (SP 8L:16D), we subjected wild type (WT) (*w*^*1118*^) flies to a reproductive dormancy assay in which they were reared for 28 days at 10°C in either LP or SP. Under SP, flies showed significantly smaller ovary size when compared to flies maintained in LP (Fig. 1A). This provides a suitable readout to investigate the effect of *eya* genetic manipulation in the photoperiodic control of seasonal physiology. As shown in previous studies (18), we found that temperature above 10-12°C was not sufficient to induce *D. melanogaster* reproductive dormancy in SP.

**Figure 1.**
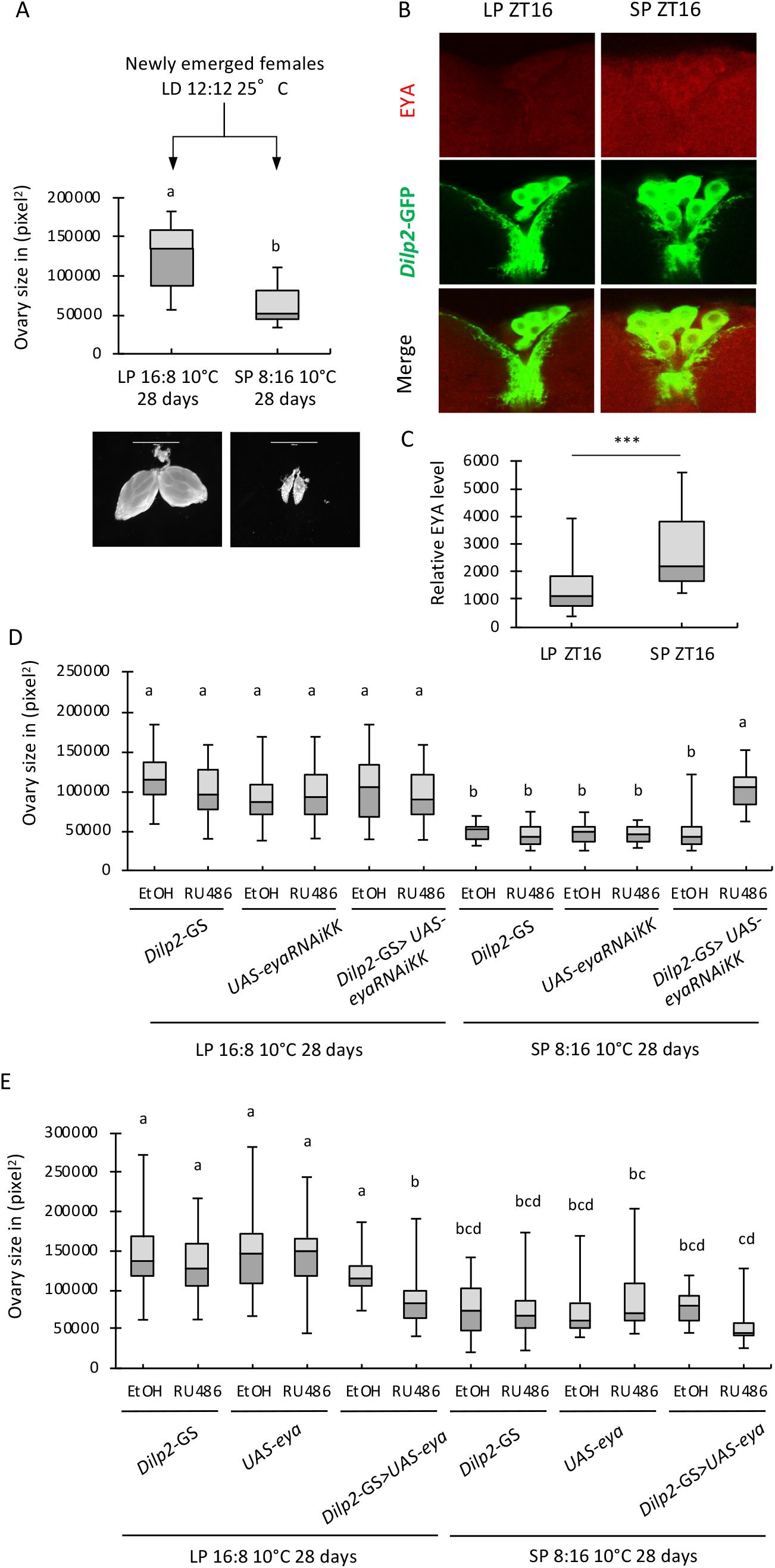
Genetic manipulation of *eya* in the *pars intercerebralis* (PI) impacts reproductive dormancy. (A) Levels of reproductive dormancy determined by ovary size measurement (in pixel^2^) in wild type (*w*^*1118*^) females reared for 28 days at 10°C in long photoperiod (LP 16L:8D) and short photoperiod (SP 8L: 16D). The whisker caps represent the minimum and maximum values, different letters indicate significant differences in ovary size between groups. All error bars indicate standard error of the mean (SEM). Two-way ANOVA, p<0.0001, n=40. (B) Comparison of EYA levels in *dilp2-gal4>UAS-cd8-GFP* brains between LP and SP at ZT16 (10°C) and (C) quantification of EYA staining in IPCs. 5-6 neurons per brain were imaged for 8-9 brains, ***p<0.001, Mann Whitney test. Ovary size of females (D) expressing *eya* dsRNAs (*dilp2-GS>eyaRNAiKK* in the presence of RU486) or (E) overexpressing *eya* (*dilp2-GS>UAS-eya* in the presence of RU486) in *dilp2* neurons at adult stage as compared to parental controls and vehicle control (EtOH) in LP and SP. Kruskall-Wallis test with Dunn’s multiple comparison test, p<0.001, n= 30 to 40 per group.

To ensure that the observed differences in reproductive dormancy is primarily driven by photoperiod length and not heavily dependent on minor thermal variations (< +/−0.4°C) associated with duration of light period in LP vs SP, we abolished temperature variations between light and dark periods by adjusting incubator temperature settings and evaluated reproductive dormancy in WT flies (Fig. S1A). In LP, WT females exhibited significantly smaller ovaries when temperature variations between light and dark periods were abolished (Fig. S1B). Most importantly WT ovaries in SP without temperature variations between light and dark periods were still significantly smaller than the corresponding ovaries in LP. While we observed an additive effect of temperature cycles in promoting larger ovaries in LP, we conclude that the difference in photoperiod length alone is sufficient to produce significant differences in ovary development at 10°C.

### *eya* mediates photoperiodic regulation of reproductive dormancy

To lay the groundwork for tissue-specific manipulation of *eya* expression, we performed immunostaining in adult fly brains by driving GFP expression under *eya* promoter. We observed GFP expression in the optic lobes at ZT8 (LD12:12 at 25°C) (Fig. S2). We also observed GFP expression in insulin producing cells (IPCs) within the *pars intercerebralis* (PI) and dorsal lateral peptidergic (DLPs) neurons located in the *pars lateralis* (PL). Expression pattern of *eya* correlates with its confirmed role in eye development as well as its potential role in regulating neuroendocrinology of seasonality. To confirm EYA expression in IPCs, we performed EYA and GFP double staining in *dilp2-Gal4>UAS-GFP* fly brains and compared EYA levels between LP and SP (ZT16) at 10°C. The choice of ZT16 was justified by EYA protein analysis described later in this study. Our results clearly indicated the presence of EYA in IPCs. In addition, we observed significantly higher EYA signal in *dilp2* expressing neurons under SP (Fig. 1B and C), consistent with the potential role of EYA in promoting reproductive dormancy under SP.

We first tested the requirement of *eya* expression in the PI for reproductive dormancy as it has been identified as an important brain region for neuroendocrine control of insect physiology. In particular, insulin signaling has been shown to be an important regulator of seasonal adaptation (23–27). *Dilp2-GS* gene-switch Gal4 driver (28) was used to drive the expression of *eya* double-stranded (ds) RNAs in the IPCs within the PI at the adult stage to knock down *eya* in a time- and tissue-specific manner to limit pleiotropic and developmental effects (Fig. 1D). Reduction in *eya* expression has no effect on ovary size in LP at 10°C but led to a significant increase in ovary size in SP at 10°C when compared to all control groups. Conversely when we overexpressed *eya* in *dilp2* expressing neurons at the adult stage, females showed significant decrease in ovary size when compared to parental controls and vehicle control in LP at 10°C (Fig. 1E). The ovaries of *eya* overexpressors in LP were comparable to that observed in flies reared in SP.

We also tested the importance of *eya* expression in *gmr*-expressing cells in the visual system (Fig. S3A and B) since we observed prominent *eya* expression in the optic lobes (Fig. S2). Knock down of *eya* using *gmr-GS* gene switch driver (29) at the adult stage similarly resulted in significant increase in ovary size specifically in SP at 10°C (Fig. S3A). On the other hand, *eya* overexpression in *gmr*-expressing cells led to significant decrease in ovary size even in LP (Fig. S3B). These observations are consistent with our results using *eya-Gal4* driver to broadly overexpress *eya* (Fig. S1C and S3C). To our knowledge, our results provide the first functional evidence supporting the involvement of *eya* in the regulation of reproductive dormancy in arthropods.

### EYA senses both photoperiod and temperature

Since photoperiodic signal constitutes a major predictable cue to anticipate seasonal progression, we hypothesize that *eya* may interpret photoperiod length and modulate its expression at either transcriptional and/or protein level to regulate seasonal physiology. To test this hypothesis, we evaluated daily *eya* mRNA and EYA protein expression from head extracts of WT (*w*^*1118*^) flies before and after application of a photoperiodic shift by rearing flies for 3 days in LP followed by 7 days in SP (Fig. 2A). Temperature was set at 10°C throughout the entire experiment after fly emergence in order to maintain conditions necessary for reproductive dormancy and specifically assess the contribution of photoperiod. Samples were collected at day 3, 5 and 10 respectively (LP3, SP5 and SP10) and assayed by qPCR and Western blotting (Fig. 2B to G). *eya* mRNA and proteins were analyzed at multiple time-points over a 24-hour light-dark cycle as mammalian *Eya3* mRNA was shown to be clock-regulated (30). Analysis of EYA proteins on LP3, SP5 and SP10 showed robust oscillation over the daily cycle in all 3 groups. A striking effect of the photoperiodic shift was an 8-hr delay in peak phase at the protein level between LP (ZT8) and SP (ZT16). This corresponds precisely to the difference in day length between the respective photoperiodic regimes. Unlike EYA proteins, the phase of *eya* mRNA oscillations remains unchanged between LP and SP conditions (Fig. 2E to G). This observation suggests a possible role of EYA in sensing day length at the protein level, specifically peaking in the dark phase in SP.

**Figure 2.**
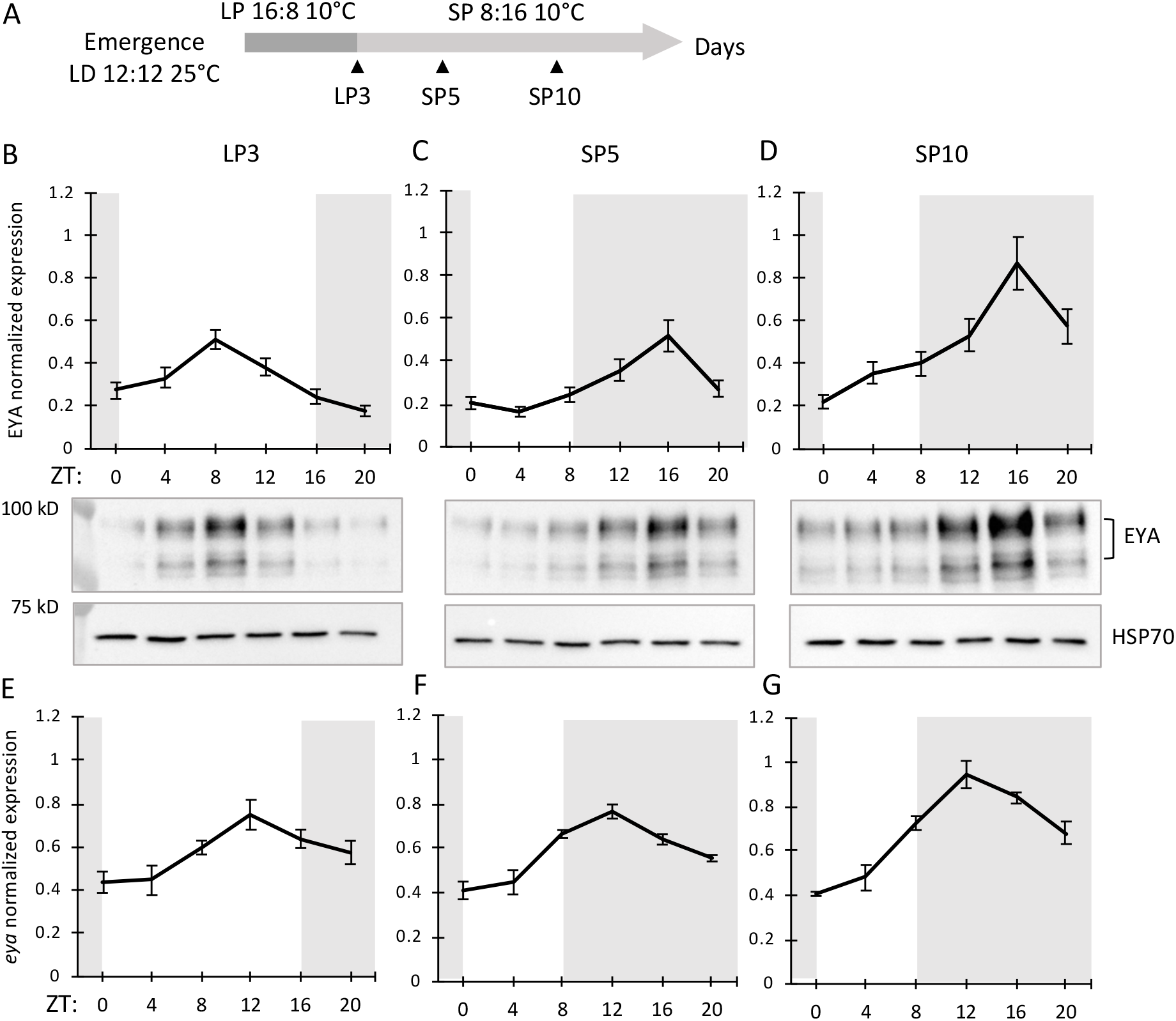
The peak phase of EY*A* protein expression is regulated by photoperiod. (A) Schematic diagram illustrating conditions and collection time-points for photoperiodic shift experiments (black arrows denote collection time-points). Samples were collected on LP3, SP5, and SP10 days at 4h intervals over 24h periods. (B, C, D) Western blots and quantifications comparing EYA expression profiles between head extracts of *w*^*1118*^ on LP3, SP5 and SP10 (peak phase: LP3=ZT8, SP5 and SP10=ZT16, p<0.01 for all conditions, JTK-CYCLE). EYA protein levels were detected using α-EYA10H6 (top isoform was used for quantification). α-HSP70 was used to indicate equal loading and for normalization. (E, F, G) Comparison of *eya* mRNA expression in heads of *w*^*1118*^ flies between LP3, SP5, and SP10 normalized to *cbp20* (peak phase: all at ZT12, p<0.05 for all conditions, JTK-CYCLE). The grey shading on each graph indicates when lights were off during each sampling period (ZT: Zeitgeber time (hr)). Data are mean ± SEM of n=2 biological replicates for mRNA and n=3 biological replicates for protein analysis. Technical duplicates were performed for each biological replicate for mRNA analysis.

Interestingly, we observed a significant increase between SP5 and SP10 in peak *eya* mRNA expression at ZT12 (1.51-fold change) and EYA protein at ZT16 (1.67-fold change) respectively (Fig. 2; p<0.05, two-way ANOVA with *post-hoc* Tukey’s HSD tests). This led to the hypothesis that a prolonged exposure of SP perhaps in combination with low temperature might be responsible for the higher level of *eya* mRNA and EYA protein on SP10. To test this hypothesis, flies were reared in light-dark cycle (LD 12L:12D) for 3 days at 25°C (3D25) before being transferred to either 10°C (10D10) or kept at 25°C (10D25) for another 7 days (Fig. 3A). As expected, we observed an increase between 10D25 and 10D10 with a 1.70- and 1.44-fold change in *eya* transcript at ZT12 (p<0.05, two-way ANOVA with *post-hoc* Tukey’s HSD tests) and EYA protein at ZT16 (p<0.001) respectively (Fig. 3B to G). This supports a model in which *eya* is a cold-induced gene in *D. melanogaster* and the cold induction of EYA protein can at least be partially attributed to induction at the transcriptional level.

**Figure 3.**
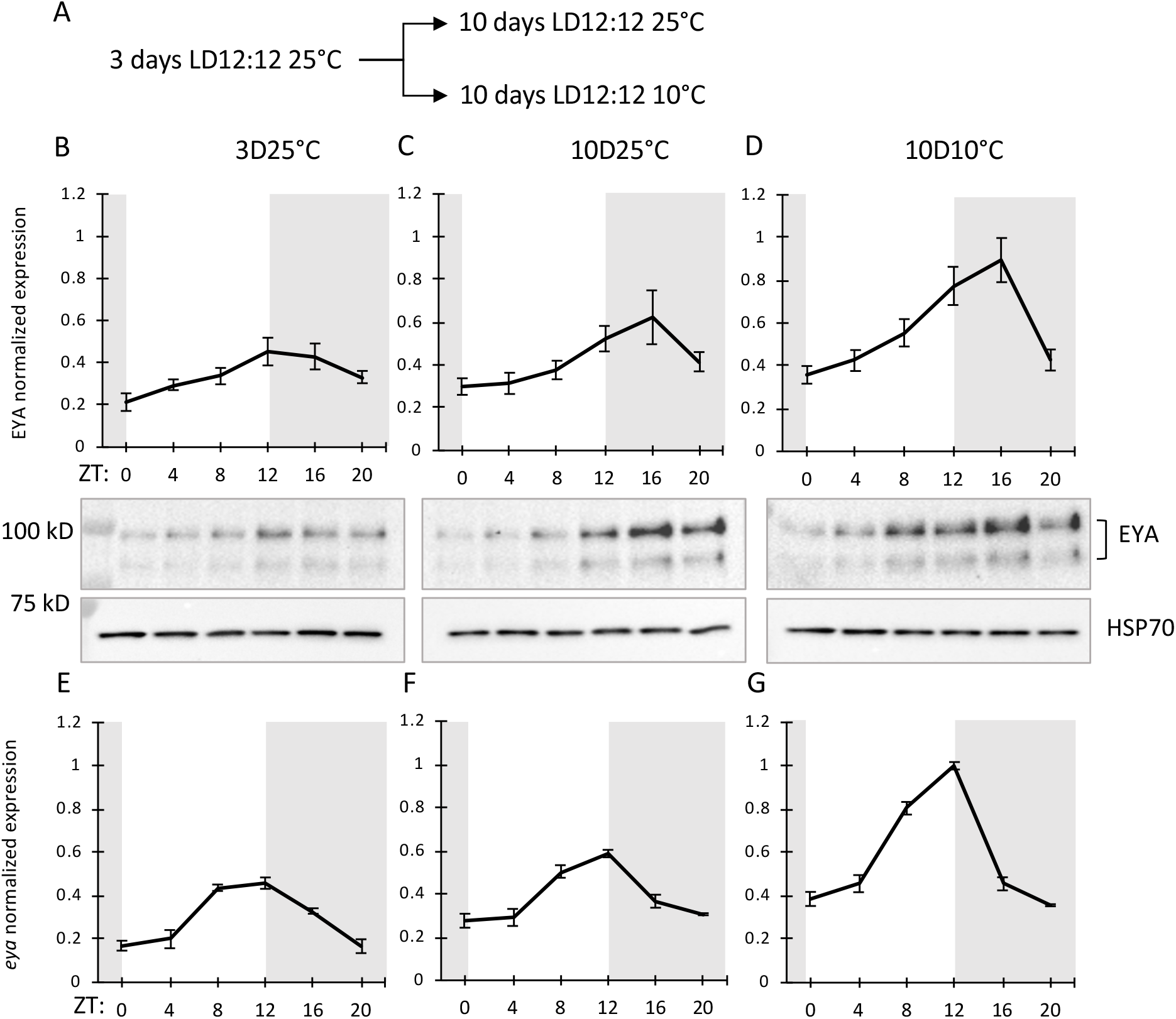
Low temperature induces significant increase in *eya* expression and amplitude at both transcriptional and protein levels. (A) Schematic diagram depicting experimental conditions for testing effect of temperature on *eya* mRNA and EYA protein. Newly emerged flies were reared at 10°C for 3 days in LD12:12 at 25°C followed by 7 days at either 25°C or 10°C. Flies were collected on day 3 prior to shifting half the flies into 10°C and on day 10 at 4hr intervals over 24hr periods. (B, C, D) Western blots and quantifications comparing EYA expression profiles between head extracts of *w*^*1118*^ on 3d 25°C, 10d 25°C and 10d 10°C (peak phase: 3D25= ZT14, 10D25=ZT16, 10D10= ZT14, p<0.0001 for all conditions, JTK-CYCLE). EYA levels were detected using α-EYA10H6 (top isoform was used for quantification). α-HSP70 was used to indicate equal loading and for normalization. (E, F, G) Comparison of *eya* mRNA expression in heads of *w*^*1118*^ flies between 3d 25°C, 10d 25°C and 10d 10°C normalized to *cbp20* (peak phase: all at ZT12, p<0.05 for all conditions, JTK-CYCLE). The grey shading in each graph indicates when lights were off during each sampling period (ZT: Zeitgeber time (hr)). Data are mean ± SEM of n=2 biological replicates for mRNA and n=3 biological replicates for protein analysis. Technical duplicates were performed for each biological replicate for mRNA analysis.

### EYA rhythmic expression is not directly controlled by the circadian clock

To determine whether *eya* daily oscillation is regulated by the circadian clock, we examined its expression in WT (*w*^*1118*^) flies in free-running condition and in *Clk* mutants (*clk*^*out*^) using qPCR. Although *eya* mRNA is rhythmic in LD cycles, these oscillations do not persist in constant darkness (DD) (Fig. 4A). This suggests that *eya* rhythmic expression may be regulated through a light-dependent pathway rather than being clock-controlled. This result is supported by the fact that no significant change in *eya* mRNA expression and oscillation was observed in *clk*^*out*^ mutants (Fig. 4B).

**Figure 4.**
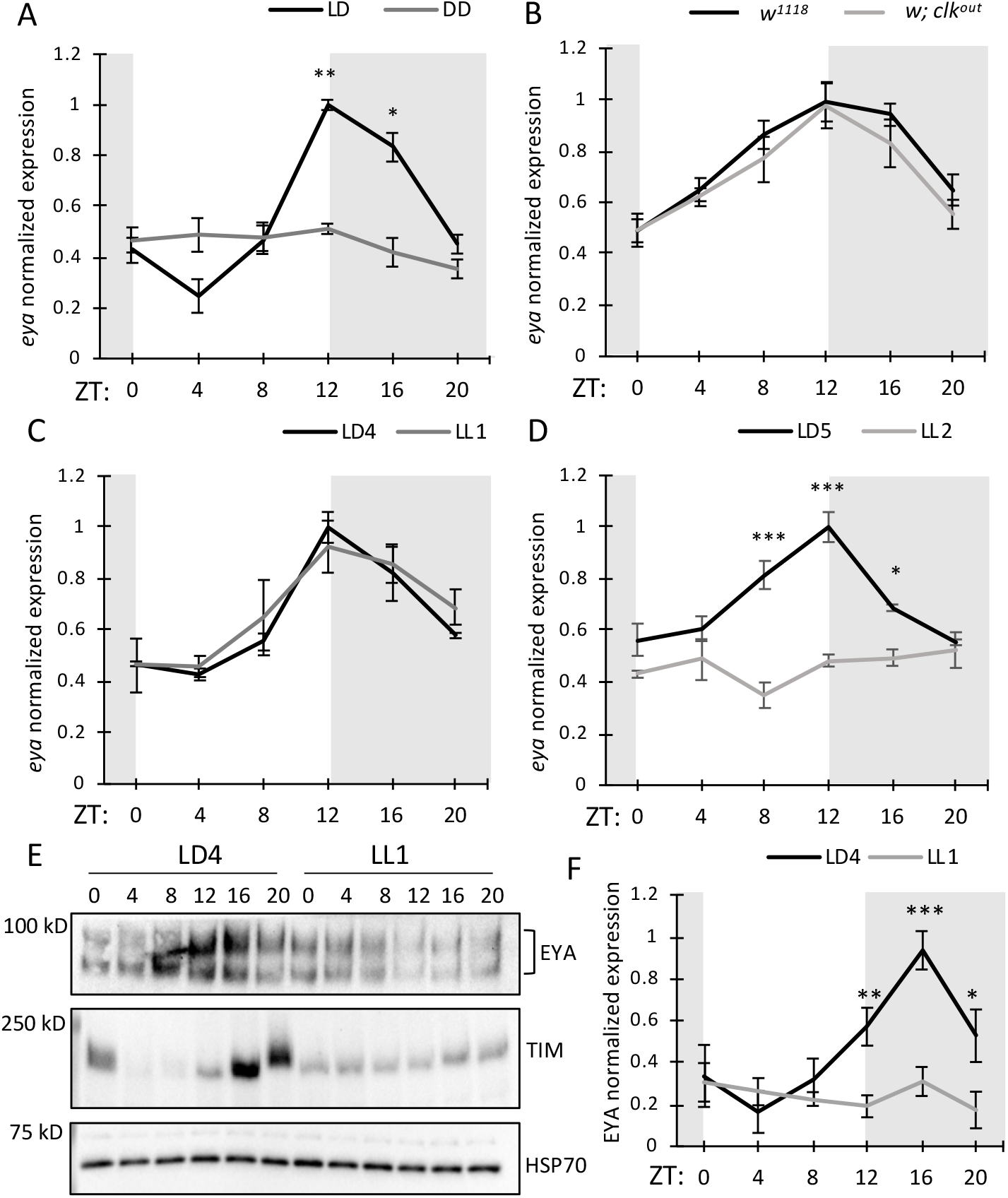
Light affects EYA protein stability. (A) *eya* mRNA expression in heads of *w*^*1118*^ flies collected on LD4 and DD1 after 3 days of entrainment (LD1-3) at LD12:12 at 25°C. (LD: peak phase=ZT12, p(LD)<0.0001; p(DD)=1, JTK-CYCLE). (B) *eya* mRNA expression in heads of *w*^*1118*^ compared to *w*; *clk*^*out*^ in LD (12:12). Flies were harvested on LD4 at LD12:12 at 25°C (peak=ZT12, p(WT)<0.0001, p(*clk*^*out*^)<0.005, JTK-CYCLE). (C, D) *eya* mRNA expression in heads of *w*^*1118*^ flies collected on (C) LD4 and LL1 (peak=ZT12, p(LD4)<0.0001, p(LL1)<0.001, JTK-CYCLE) and on (D) LD5 and LL2 (peak=ZT12, p(LD5)<0.0001 ; p(LL2)=1, JTK-CYCLE) after 3 days of entrainment at LD12:12 at 25°C. (E) Western blots comparing EYA expression profiles in heads of *w*^*1118*^ flies collected on LD4 and LL1. (F) Quantification of (E) and expressed as relative expression (highest value = 1) (peak=ZT16, p(LD4)<0.005; p(LL1)=1, JTK-CYCLE). Data are mean ± SEM of n=2 biological replicates. Technical duplicates were performed for each biological replicate for mRNA analysis. Asterisks denote significant differences between conditions or genotypes at each ZT: ***p<0.001, **p<0.01, *p<0.05, two-way ANOVA with *post-hoc* Tukey’s HSD tests.

To further examine the role of light in the regulation of *eya* mRNA, we compared expression of *eya* in WT flies in LD to those in constant light condition (LL1 to LL2) (Fig. 4C and D). *eya* daily mRNA expression remains unaffected in LL1 compared to LD4 after 3 days of LD entrainment (Fig. 4C). However when comparing LD5 vs LL2, *eya* mRNA abundance was significantly reduced in LL2 at ZT8, 12, and 16 and no oscillation was detected. Interestingly, EYA protein level was significantly reduced even on LL1 as compared to LD4 and the oscillation was dampened (Fig. 4E and F). The effect of light on EYA protein stability shows striking similarities with the well-characterized light-mediated TIM degradation (31–33) (Fig. 4E, middle panel). One difference between EYA and TIM expression in LD is that EYA appears to accumulate at earlier time-points compared to TIM, which shows an expression pattern restricted to the dark phase only.

### Genetic manipulation of *tim* alters EYA stability and reproductive dormancy

Since EYA protein exhibits a similar pattern as TIM protein under LL, we decided to examine its expression level in *tim* mutant flies to investigate whether TIM is involved in stabilizing EYA. Genetic variations in *tim* has previously been linked to latitudinal clines in diapause incidence in *Drosophila* (34, 35). By Western blotting, we observed that EYA abundance is significantly reduced in *tim* null mutants (*yw*; *tim*^*0*^) from ZT4 to ZT16 (Fig. 5A and B) and starts to accumulate after lights-off. Conversely, a significant increase in EYA was detected in *tim*-overexpressing flies (*w*; *p*{*tim(WT)*}) from ZT4 to ZT20 (Fig. 5C and D). Interestingly, no significant change in *eya* mRNA expression was detected when comparing WT to *tim* null mutants or to *tim* overexpressors at all time-points (Fig. 5E and F), supporting the role of TIM in stabilizing EYA at the protein level.

**Figure 5.**
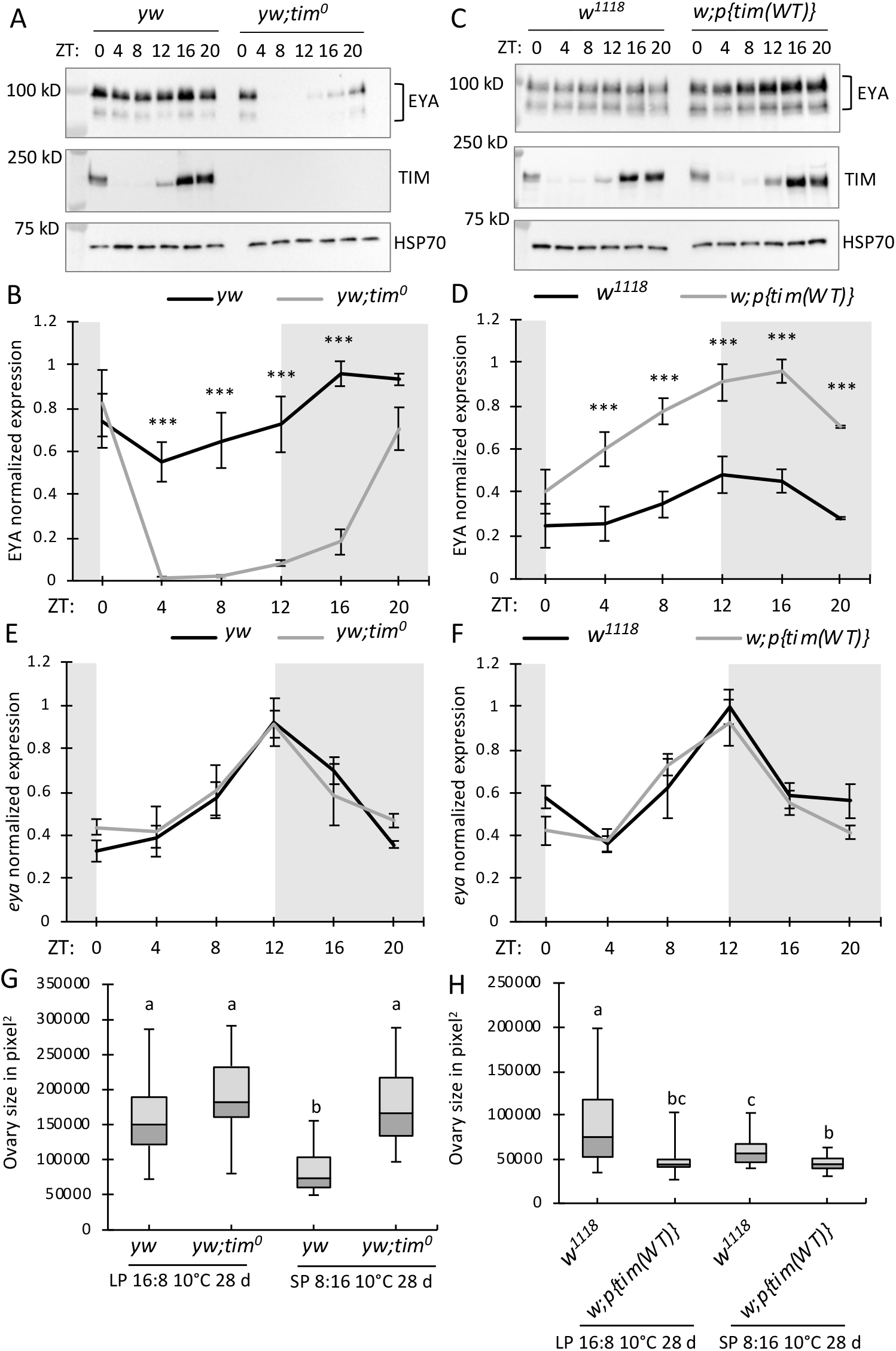
Genetic manipulation of *tim* affects EYA protein stability and incidence of reproductive dormancy. (A, C) Western blots comparing EYA expression profiles in *tim* null mutant (*yw*; *tim*^*0*^) and *tim* overexpressor (*w*^*1118*^; *p{tim(WT)})* to WT (*yw* or *w*^*1118*^) controls. Flies were entrained in LD (12:12) at 25°C for 3 days and collected at 6 time-points on LD4. (B, D) Quantification of western blots shown in (A, C) and expressed as relative expression (highest value = 1). (E, F) Daily *eya* mRNA expression in *tim* null mutant (*yw*; *tim*^*0*^) and *tim* overexpressor (*w*^*1118*^; *p{tim(WT)})* as compared to respective WT controls. Data are mean ± SEM of n=2 biological replicates. Technical duplicates were performed for each biological replicate for mRNA analysis. Asterisks denote significant differences between conditions at each ZT: ***p<0.001, **p<0.01, *p<0.05, two-way ANOVA with *post-hoc* Tukey’s HSD tests. (G, H) Ovary size of female *tim* null (*yw*; *tim*^*0*^) and *tim* overexpressor (*w*; *p{tim(WT)})* as compared to WT (*yw* or *w*^*1118*^) in reproductive dormancy assay. The whisker caps represent the minimum and maximum values, different letters indicate significant differences in ovary size between groups with p<0.001, except for difference between b and c in (H) p<0.05, n=40 females per group, one-way ANOVA followed by *post-hoc* Tukey test.

We then proceeded to test the effect of *tim* genetic manipulations on female reproductive dormancy. We found that when *tim* null mutants were reared in SP at 10°C, their ability to enter into reproductive dormancy was significantly impaired as compared to WT (Fig. 5G). In a reciprocal experiment, we evaluated ovary size of *tim*-overexpressing flies in LP and SP and observed that the proportion of individuals in reproductive dormancy was significantly higher as compared to WT even in LP (Fig. 5H). The difference in ovary size between WT and *tim*-overexpressing flies were also significant in SP but much smaller (Fig. 5H). Our results are consistent with the role of TIM in supporting EYA in the regulation of reproductive dormancy in response to photoperiodic signals.

### TIM mediates both photoperiodic and temperature signals to impact EYA expression

Although analysis of *tim* mutants support the hypothesis that TIM may promote EYA expression, the low level of canonical TIM expression at low temperature (36) and its absence during the light phase (31–33) suggest that other TIM isoforms could be involved. We therefore evaluated whether the expression profiles of different TIM isoforms are affected by temperature shift in a similar fashion as EYA. At 10°C, canonical TIM was barely detectable using our TIM antibody when compared to TIM levels at 25°C and no other isoforms were detected. Concomitantly, recent studies revealed that *tim* undergoes a thermosensitive alternative splicing controlling the relative abundance of various *tim* isoforms (37, 38). Consistent with our observation, the authors found lower levels of the canonical TIM protein at 18°C and revealed the induction of two cold-specific splice forms (*tim-cold and tim-short&cold*). The antibody we used to detect TIM was generated using a C-terminal antigen that is absent in the shorter TIM-SC isoform. In order to detect TIM-SC, we used a TIM antibody generated against a N-terminal antigen (39) and repeated the analysis.

This time, we observed a drastic switch between the canonical TIM-L and the cold-induced TIM-SC isoform between 25°C and 10°C (Fig. 6A). TIM-L abundance and its cycling amplitude were significantly reduced at 10°C as compared to that observed at 25°C (Fig. 6B). Conversely, TIM-SC isoforms showed an opposite trend with a significant increase in expression at 10°C whereas almost no signal was detected at 25°C (Fig. 6C). TIM-SC was expressed at high levels at 10°C and exhibited a weak oscillation, however this isoform failed to reach significance for JTK cycling statistics likely due to its stability in the presence of light.

**Figure 6.**
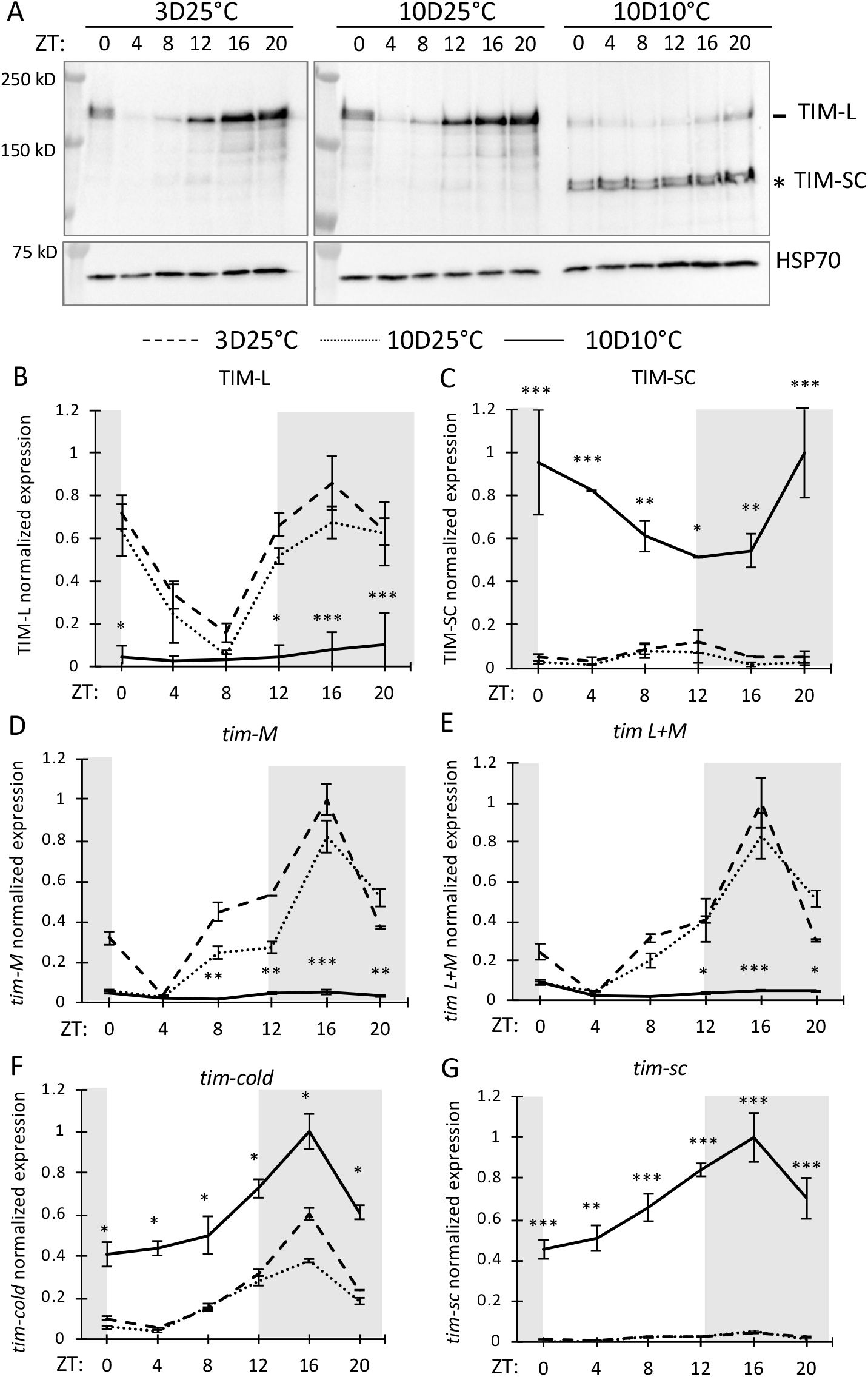
Expression of TIM-L and TIM-SC is temperature-dependent. Flies were reared using the same condition as in Figure 3. (A) Western blots comparing TIM expression profiles between head extracts of *w*^*1118*^ collected on 3d 25°C, 10d 25°C, and 10d 10°C. Both TIM-L and TIM-SC isoforms were detected using α-TIM (a gift from M. Young’s lab) (39, 69) and α-HSP70 was used to indicate equal loading and for normalization. (B, C) Quantification of western blot signals shown in (A) and expressed as relative expression (highest value = 1). (B) peak=ZT16, p(3D25)<0.0001, p(10D25)<0.0001; p(10D10)=1, JTK-CYCLE. (C) p(3D25)=1, p(10D25)=1, p(10D10)=0.064, JTK-CYCLE. (D, E, F, G) Comparison of *tim*-*M*, *tim-L+M*, *tim-cold and tim-sc* mRNA expression profiles in heads of *w*^*1118*^ flies between 3d 25°C, 10d 25°C, and 10d 10°C normalized to *cbp20*. Data are mean ± SEM of n=2 biological replicates. Technical duplicates were performed for each biological replicate for mRNA analysis. Asterisks denote significant differences between 10d 25°C and 10d 10°C at each ZT: ***p<0.001, **p<0.01, *p<0.05, two-way ANOVA with *post-hoc* Tukey’s HSD tests.

The evaluation of the mRNA levels for different *tim* isoforms confirmed that both *tim*-*cold* and *tim-sc* are induced at 10°C compared to 25°C (Fig. 6F and G), while *tim-M* and *tim-L+M* are drastically reduced in the cold (Fig. 6D and E). All together, these results support the hypothesis that *tim* thermosensitive alternative splicing plays an important role in stabilizing EYA in cold temperature.

We also investigated whether expression of different *tim* isoforms are modulated by photoperiodic signals at 10°C. At the transcriptional level, we observed a significant elevation in *tim-sc* mRNA between ZT16 and ZT20 in response to photoperiodic shift from LP to SP at 10°C (LP3 vs SP10) whereas no significant changes were detected for the longer *tim* isoforms (*tim-L+M*, *tim-M* and *tim*-*cold*) (Fig. S4A to D). In addition, we observed a 4-hr phase advance in *tim*-*sc* expression peak as flies transitioned from LP to SP condition (Fig. S4D). At the protein level, consistent with the effect of low temperature on canonical TIM protein, we observed relatively low level of TIM-L across LP and SP conditions at 10°C (Fig. S4E). In SP5 and even more so in SP10, significantly higher levels of TIM-L were observed at ZT0 and from ZT12/16 to ZT20 as compared to TIM-L at LP3 (Fig. S4F). Robust daily cycling of TIM-L was detected in SP5 and SP10 at 10°C, but not in LP3. Despite the effect of low temperature in diminishing TIM abundance, SP may still play a role in increasing TIM-L accumulation during night time, likely due to increased duration of the dark phase. With regard to TIM-SC, no change in peak phase was observed between LP and SP although we detected significantly higher level in TIM-SC at ZT20 in SP10 as compared to LP3 (Fig. S4G). Combining the results of temperature and photoperiodic shift experiments, SP in conjunction with low temperature provide the ideal condition for higher TIM-L and TIM-SC isoforms at night as well as elevated abundance of TIM-SC during the day. Both isoforms may function to stabilize EYA and promote reproductive dormancy. This is consistent with more abundant EYA and increased incidence of reproductive dormancy in flies overexpressing *tim*.

Finally to test the potential of TIM in stabilizing EYA through protein interactions, we evaluated the ability of TIM-L and TIM-SC isoforms to bind EYA by expressing them in *Drosophila* S2 cells. Reciprocal coimmunoprecipitations suggested that both TIM isoforms can interact with EYA (Fig. 7A), although the affinity of TIM-SC to EYA is significantly higher as compared to TIM-L (Fig. 7B). To test whether EYA and TIM could colocalize in specific brain regions involved in photoperiodic sensing, we expressed GFP under the control of *eya* promoter and stained for TIM using the antibody that recognizes both TIM-L and TIM-SC isoforms. While we cannot rule out the possibility that we are observing some expression for TIM-L, it is reasonable to believe that most of the TIM signal observed corresponds to TIM-SC expression at 10°C. We performed our analysis at ZT16 on SP5 since TIM and EYA expression was shown to be abundant by Western blot analysis. Analysis of whole brain immunostaining revealed colocalization of *eya* and TIM in the optic lobes. While we detected signal for both *eya* and TIM in the PI, we did not find evidence for colocalization (Fig. 7C to E).

**Figure 7.**
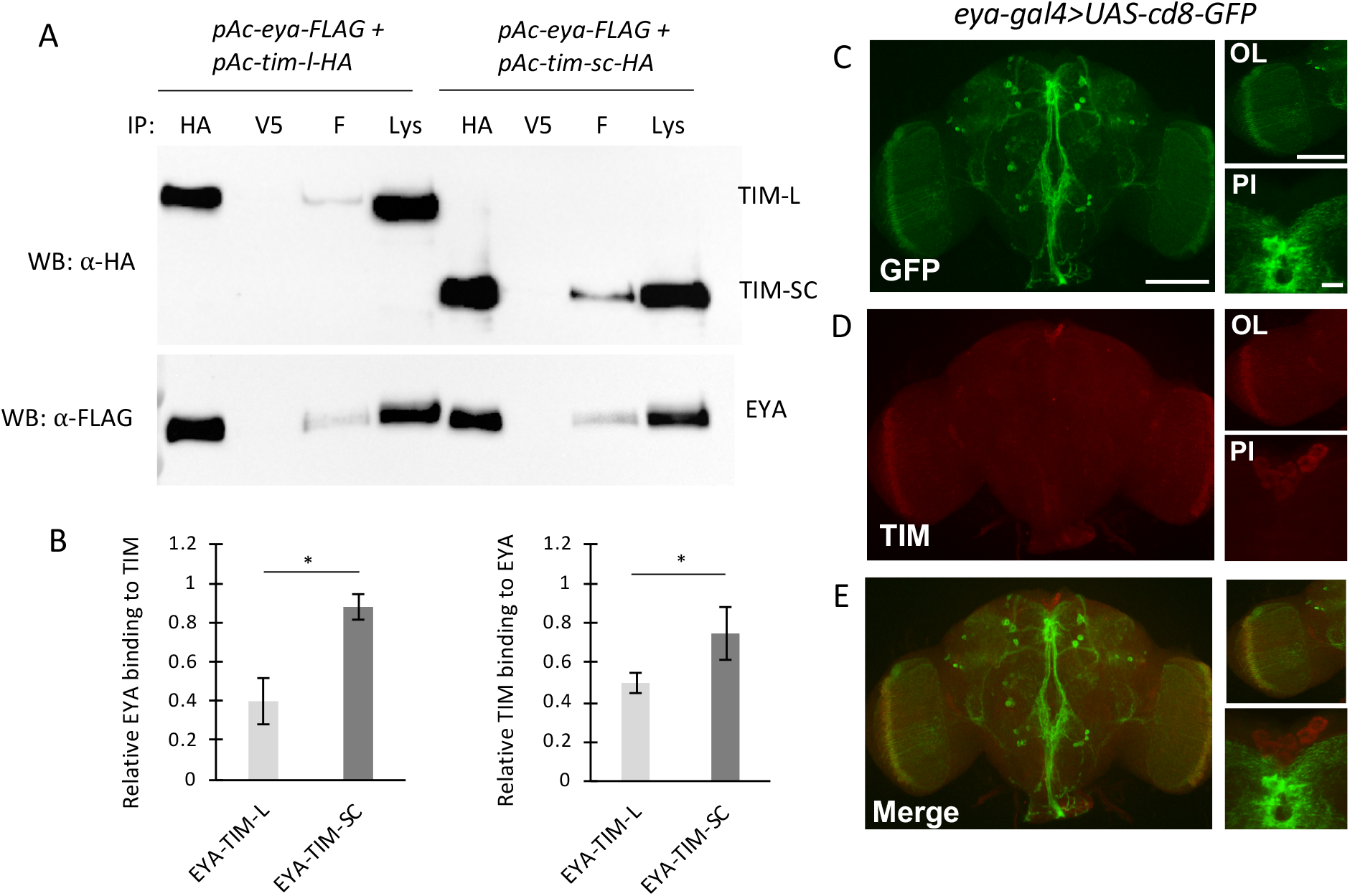
TIM-SC shows higher affinity for EYA as compared to TIM-L. (A) Western blots showing results of reciprocal coimmunoprecipitation (coIP) assays to detect interaction of EYA with TIM-L and TIM-SC in *Drosophila* S2 cells. Proteins extracted from S2 cells were either immunoprecipitated with α-FLAG to pull down EYA or with α-HA to pull down either TIM-L or TIM-SC. Negative control coIPs were performed using α-V5, which do not recognize the proteins of interest. Immuno-complexes were subjected to western blotting to detect the bait protein or protein interactions. Input for the coIP is indicated (Lys). (B) Bar graphs showing quantification of reciprocal coIP assays (n=4). Values for binding of interacting proteins were normalized to amount of bait detected in the respective IPs. Error bars indicate +/−SEM (* p<0.05, two-tailed Student’s t test). (C-E) *eya-gal4/UAS-cd8-GFP* brains from flies collected on SP5 (ZT16) at 10°C were stained for (C) GFP and (D) TIM. PI: *pars intercerebralis*; OL: optic lobes (OL). (E) Merged images showing colocalization of GFP (*eya*) and TIM in the OL but not in the PI. 8-9 brains were imaged.

## Discussion

In this study we provide converging evidence supporting a central role of *eya* in seasonal adaptation in *D. melanogaster* (Fig. S5). This includes immunocytochemical detection of EYA in areas within the fly brain regulating diapause and insulin signaling and the capacity of *eya* mRNA and protein to track temperature and photoperiod shifts through modulation of their expression. Perhaps the strongest evidence is our finding that spatiotemporal genetic manipulation of *eya* can alter seasonal physiology in the context of female reproductive dormancy. Our results support a conserved role of *eya* in regulating seasonal physiology in response to environmental cues in animals, but also highlight key differences between the mechanisms by which *eya* and *Eya3* perform their functions in *D. melanogaster* and sheep.

In *Drosophila*, *eya* was first characterized as a key player for eye development and subsequently more broadly for organ development (8, 40). Our analysis of *eya* expression in the adult fly brain provides strong indication that *eya* functions beyond development. We observed that *eya* is expressed in specific regions within the brain: the PI and the PL, which house IPCs and DLPs neurons, respectively (41). Interestingly these two clusters of neurosecretory cells play an important role in the hormonal control of insect diapause (25, 42). In a large number of insect species, it is well established that endocrine regulation of diapause in adult females is mediated by ILPs and Juvenile Hormone (JH) through the inactivation of the IPCs/CA/ovarian axis (23–27, 43, 44). In *D. melanogaster*, decreased IPC activity is associated with downregulation of *ilp2* and *ilp3* production and correlates with an increased incidence of reproductive dormancy (26). It is possible that the function of EYA in photoperiodic sensing is at least partially mediated through insulin signaling.

In addition to neurosecretory cells, we also observed *eya* expression in adult optic lobes. Previous work demonstrated that the absence of compound eyes in *cli*^*eya*^ mutants affect their ability to entrain to extremely short or long photoperiods (45). The fact that knock down of *eya* in adult visual system can disrupt photoperiodic sensing indicates that EYA functions in transmitting photoperiodic signals in addition to its role in compound eye development. Compound eyes are necessary for phase adjustment and considering the important role of the visual system in light transduction (46), it is possible that EYA might play a role in mediating photoperiodic signals to other sites within the brain.

Although the use of *D. melanogaster* as a model to study insect photoperiodic timing has been historically challenging, there is accumulating evidence showing that adult reproductive dormancy in *D. melanogaster* can be induced by short photoperiod at sufficiently low temperature. A number of studies found that flies entrained in short photoperiod (<14 hr of light per day) and low temperature (<12°C) show significantly reduced vitellogenesis compared to females exposed to long photoperiod (16 hr of light per day) at the same low temperature (44, 47, 48). In synergy with light, temperature plays a major role in *Drosophila* reproductive dormancy. Indeed, regardless of the photoperiodic conditions, flies entrained at 12°C and above always undergo ovarian development. This is in contrary to other insect species for which photoperiod plays a dominant role in the induction of seasonal phenotypes (49, 50). A key step of this study was therefore to establish a robust phenotypic readout for seasonal physiology. By rearing flies at 10°C for 28 days after emergence either in LP and SP, we were able to induce significant phenotypic differences in reproductive dormancy and used these conditions to evaluate the impact of manipulating *eya* expression in physiologically relevant conditions. We found that manipulation of *eya* in *eya*-expressing cells and more specifically either in the visual system (*gmr*-expressing cells) or in insulin-producing neurons, had significant consequences on reproductive dormancy. Our results represent the first *in vivo* genetic manipulation data supporting a critical role of *eya* as a photoperiodic sensor in animals.

Another advantage of establishing robust conditions to induce reproductive dormancy in flies was the ability to track *eya* mRNA and protein expression in physiologically relevant conditions and investigate whether *eya* can “sense” photoperiodic and temperature shifts that ultimately result in different outcomes in reproductive dormancy. The mammalian ortholog *Eya3* has been proposed to play an important role in photoperiodism (11, 14, 15). In sheep, *Eya3* is a clock-controlled gene whose rhythmic expression is set by the phase of evening melatonin onset. This represents a classic example of external coincidence where light-dependent stimulus interacts with circadian oscillator to trigger seasonal phenotype (51). We asked the question whether *eya* mRNA or protein can similarly interpret photoperiodic cues in insects. We observed that the phase of EYA oscillation is strongly influenced by photoperiod while that for *eya* mRNA is not affected. Indeed in LP, EYA peak expression is detected during the day time and transferring the flies into SP induces an 8-hr shift in EYA peak phase, restricting its accumulation during the night time. In contrast, Dardente et al. observed *Eya3* photoperiodic shift occurring at the transcriptional level with an accumulation during day time in LP, consistent with *Eya3*-dependent induction of a summer phenotype in sheep (11).

In mammals, the effect of changing day length on endocrine rhythm is mediated by nocturnal secretion of melatonin by suppressing the expression of a range of E box-controlled gene, including *Eya3* (52). Melatonin has been implicated in photoperiodism in some insects species (53, 54); however, its involvement in *Drosophila* is largely debated. This could explain why we did not observe any significant variation in *eya* mRNA oscillation in response to changing photoperiods.

How *eya* is regulated at the transcriptional level in flies has not been resolved in this study. Despite the presence of E-boxes upstream of *eya* coding sequence (Fig. S6), *eya* expression was unaltered in *clk*^*out*^ mutant. In addition, daily cycling of *eya* mRNA was abolished in free-running condition, suggesting that *eya* transcription is not clock-controlled. This is consistent with recent findings suggesting that photoperiodic sensing appears independent of robust circadian clocks (55). Among the potential pathways involved in regulating *eya* transcription, Pigment Dispersing Factor (PDF) signaling represents a strong candidate. PDF is well characterized for its role in photoperiodic sensitivity (56–62) and there is accumulating evidence supporting its involvement in reproductive dormancy by conveying day length signal to the PI (63, 64). We speculate that PDF could regulate *eya* expression in a cAMP-dependent manner, perhaps in response to light signal. This is supported by the presence of CREB element in the *eya* promoter (Fig. S6). Similarly in mammals, *Eya3* transcription is regulated by cAMP (14).

Independent of the effect of light on *eya* mRNA expression, the abundance of EYA protein is significantly reduced under constant light, similar to the effect of light on TIM. To test our hypothesis that the light response of EYA is mediated by the light sensitivity of TIM and TIM may stabilize EYA at the protein level, we evaluated EYA expression in *tim* mutants. As anticipated, we observed a correlation between EYA and TIM proteins with significantly higher EYA when TIM is abundant and vice versa. In particular, under photoperiodic shift we found that all TIM isoforms (TIM-L/TIM-Cold and TIM-SC) accumulate during the night time in SP10 matching the increase in EYA levels from ZT12 to ZT16 in the same conditions. Since this experiment was conducted at 10°C, it is likely that the residual expression of the long TIM isoforms (TIM-L/TIM-Cold) we detected is in most part due to the induction of the TIM-Cold isoform. Nonetheless, due to their similarity in size, TIM-L and TIM-Cold could not be clearly differentiated by western blot.

Synergism between SP and low temperature is key to trigger appropriate seasonal response in *D. melanogaster*. Here we found that low temperature has an enhancing effect on *eya* amplitude at both transcriptional and protein levels. This agrees with the transcriptomic analysis of a recent publication that reveals an upregulation of *eya* in flies exposed to short photoperiod (10 L:14D) at 11°C for 3 weeks when compared to reproductively active condition (25°C, 12 L:12D) (65). In light of the recent evidence on *tim* thermosensitive alternative splicing (37, 38, 66), we were able to confirm the induction of the *tim-sc* mRNA and protein at 10°C. Consistent with our model, our results support the prevalent role of the cold-induced TIM-SC isoforms in stabilizing EYA under low temperature and suggest that TIM thermosensitive splicing mechanism plays a role in regulating seasonal physiology. The ability of TIM-SC to stabilize EYA appears to be due to its increased light-insensitivity and affinity to EYA as compared to canonical TIM isoforms.

The model of TIM and EYA collaboration in photoperiodic time measurement is strongly supported by the effect of *tim* genetic manipulation on reproductive dormancy. Indeed, *tim*^*0*^ flies in SP have significantly larger ovaries compared to control individuals whereas flies overexpressing *tim* undergo reproductive dormancy even in LP. All together these results suggest that the effect of light on EYA protein stability is mediated through TIM. If TIM indeed stabilizes EYA to promote reproductive dormancy, an obvious question would be the nature of the cell types in which this interaction takes place. We attempted to address this question and showed that TIM-EYA interactions primarily take place in the optic lobes, which has been shown to transmit photic signals to the circadian clock neuronal network, and subsequently to the PI. The fact that we did not observe TIM-EYA colocalization in the PI suggest that the mechanisms by which *eya* regulates seasonal physiology may differ in these two cell types.

Finally, the involvement of TIM in reproductive dormancy is supported by the positive correlation between diapause incidence and latitude associated with the higher allele frequency in *ls-tim* compared to *s-tim* observed in Northern European *Drosophila* populations (34). Interaction between CRY and ls-TIM is weaker than with s-TIM resulting in a reduced CRY-dependent TIM degradation (35, 67). All strains used in this study were genotyped in order to identify the *tim* allele present in their genome (Table S1). With the exception of *tim* overexpressing flies (*w*; *p*{*tim(WT)-3XFLAG}*), no correlation was found between *tim* allelic variation and reproductive dormancy in other strains. This indicate that the effect of *eya* genetic manipulation and *tim*^0^ on reproductive dormancy could not be attributed to the genetic background for the *tim* locus in those flies.

In summary, we propose that the role of EYA in seasonal adaptation in flies could rely on a combination of both internal and external coincidence model through its interaction with TIM where short photoperiod restricts EYA peak to the night time. The reliance of cold temperature by *D. melanogaster* to induce reproductive dormancy, a feature not necessarily shared by other insects, can be attributed to the cold induction of *eya* mRNA as well as EYA protein via TIM leading to increase in EYA above a specific threshold. This study provides insights to further decipher the complex mechanisms by which the circadian clock or clock-regulated proteins could inform the photoperiodic timer and allow organisms to anticipate environmental changes as seasons progress.

## Material and methods

Detailed materials and methods are provided in SI Materials and Methods.

### Fly stocks

UAS/Gal4 lines used for immunocytochemistry (ICC) and *eya* genetic manipulation as well as generation of *tim* overexpressor are described in SI.

### Reproductive dormancy assay

Reproductive dormancy assays are described in SI.

### Whole-Mount Brain Immunocytochemistry

Fly entrainment conditions and ICC procedures are described in SI.

### Western blotting and antibodies

Protein extraction from fly heads and western blotting procedures were performed as described in (68). Antibodies and dilutions are described in SI.

### Analysis of circadian/daily gene expression by real-time PCR

RNA extraction was performed as previously described (68). Primers used and Real-time PCR procedure are described in SI.

## Acknowledgements

We thank Patrick Emery for providing *p{tim(WT)-luc}* transgene, Michael Young/Deniz Top for providing α-TIM generated from N-terminal antigen, and Heinrich Jasper for providing *dilp2-GS* driver. We thank the Bloomington *Drosophila* Stock Center and Vienna *Drosophila* Resource Center for providing fly stocks and Developmental Studies Hybridoma Bank for providing EYA antibodies. Research in the laboratory of JCC is supported by NIH R01 GM102225, NSF IOS 1456297, and USDA NIFA SCRI 63513.

## Supplemental Information

### SI Figure Legends

**Figure S1.**
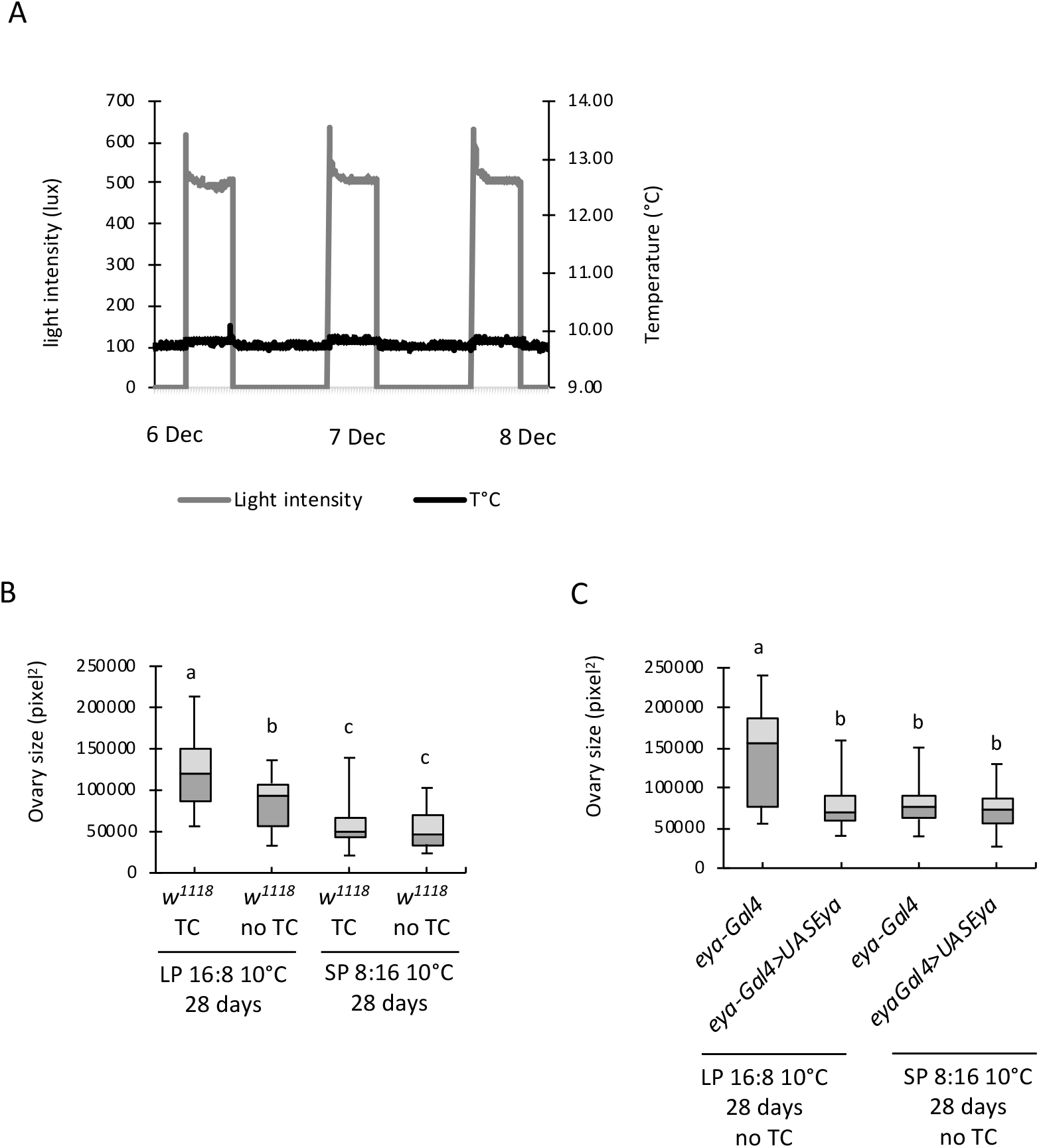
Removing minor thermal fluctuations in incubators during light vs dark phase did not abolish significant differences in ovary size between flies reared in LP and SP. (A) Temperature and light intensity recordings in incubators without temperature fluctuations between light and dark phase. (B) Ovary size (in pixel^2^) of WT females (*w*^*1118*^) reared for 28 days at 10°C in long photoperiod (LP 16L:8D) or short photoperiod (SP 8L: 16D) with or without temperature cycles (TC) or fluctuations. (C) Ovary size of females overexpressing *eya* (*eya-Gal4>UAS-eya*) as compared to parental control without temperature fluctuations between light and dark phase (no TC). The whisker caps represent the minimum and maximum values, different letters indicate significant differences in ovary size between groups with p<0.05, n=50 females per group, one-way ANOVA followed by *post-hoc* Tukey test.

**Figure S2.**
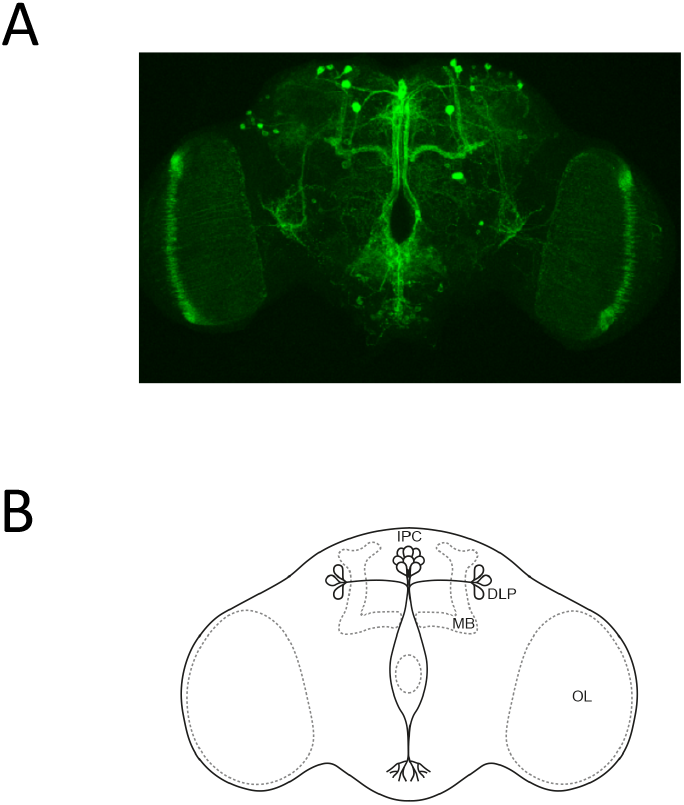
(A) *eya-gal4/UAS-cd8-GFP* brain from flies entrained in LD12:12 25°C for 3 days and dissected at ZT8 showing GFP signal in the PI region and optic lobes. (B) Schematic of *Drosophila* brain showing regions of interest OL: optic lobe, IPC: insulin producing cells, DLP: dorsal lateral peptidergic neurons, MB: mushroom bodies.

**Figure S3.**
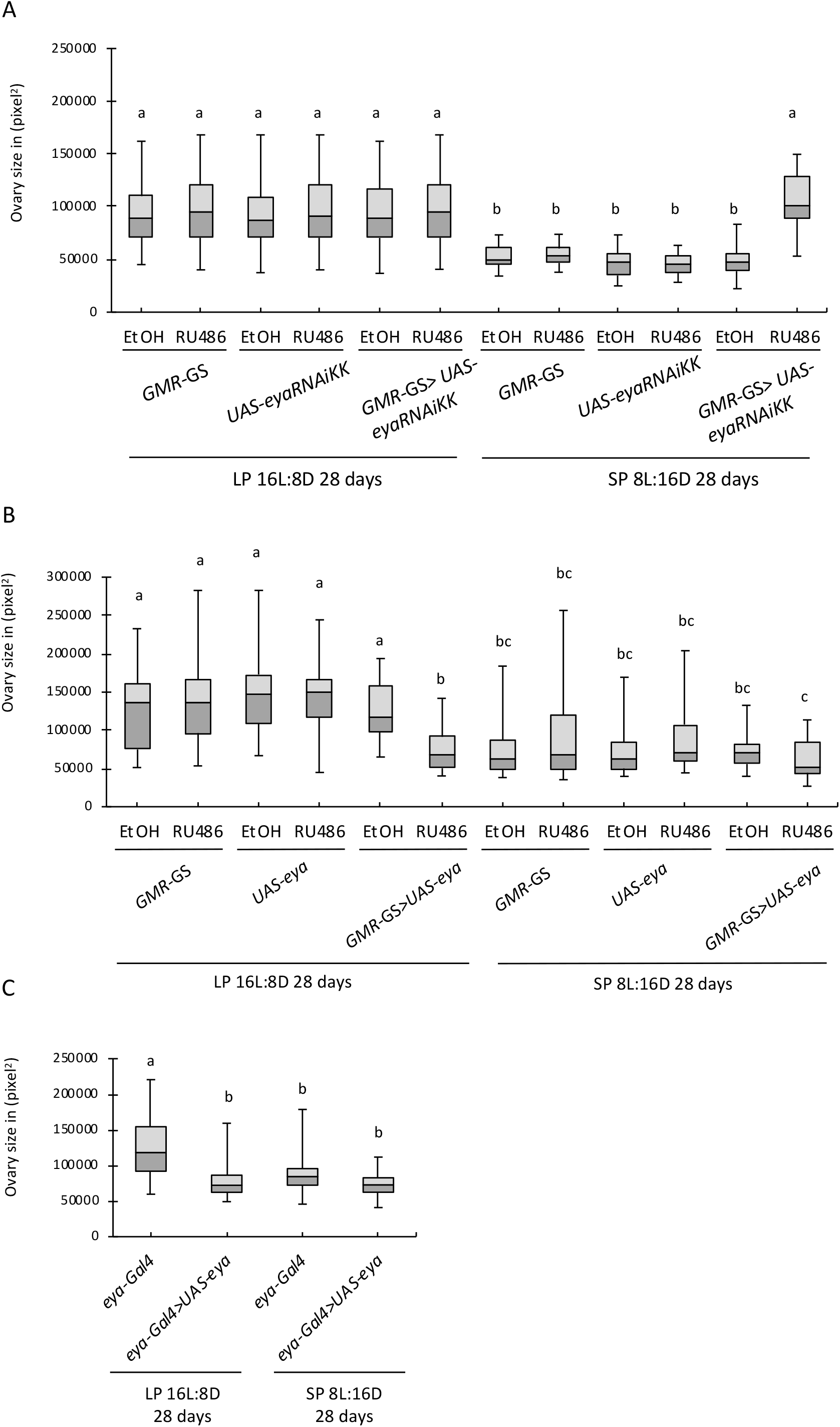
Genetic manipulation of *eya* alters incidence of reproductive dormancy. Ovary size of females (A) expressing *eya* dsRNA or (B) overexpressing *eya* in the visual system at the adult stage (*gmr-GS>eyaRNAiKK* in presence of RU486) as compared to parental controls (*gmr-GS*, *eyaRNAiKK* and *UAS-eya*). (C) Ovary size of females overexpressing *eya* in *eya*-expressing cells (*eya-Gal4>UAS-eya*) as compared to parental control. (A-C) The whisker caps represent the minimum and maximum values, different letters indicate significant differences in ovary size between groups with p<0.0001, n=30 to 40 females per group. (A, B) Kruskall-Wallis test with Dunn’s multiple comparison; (C) One-way ANOVA followed by *post-hoc* Tukey test.

**Figure S4.**
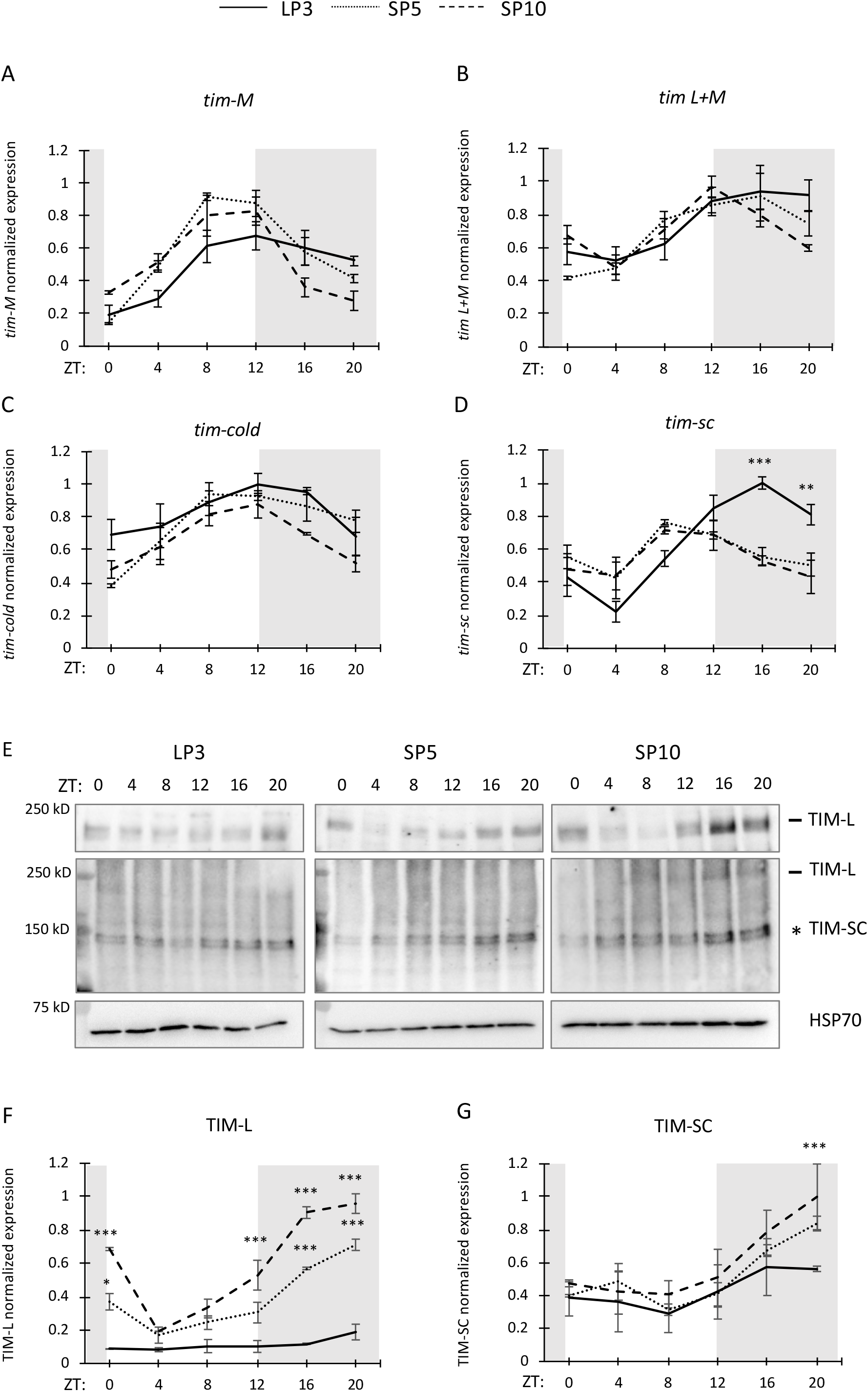
*tim* isoforms are differentially regulated in long vs short photoperiodic conditions. Flies were reared using the same condition as in Figure 2. (A, B, C, D) Comparison of *tim-M*, *tim-L+M*, *tim-cold and tim-sc* mRNA expression in heads of *w*^*1118*^ flies between LP3, SP5, and SP10 normalized to *cbp20*. Asterisks denote significant differences between LP3 and SP10 at each ZT: ***p<0.001, **p<0.01, *p<0.05, two-way ANOVA with *post-hoc* Tukey’s HSD tests. (E) Western blots comparing TIM expression profiles between head extracts of *w*^*1118*^ collected on LP3, SP5, and SP10. α-TIM C-terminus antibody (Chiu lab, RRID:AB_2782953) (top panel) and α-TIM N-terminus antibody (kindly provided by M. Young’s lab) (1, 2) (middle panel) was used to detect TIM-L and TIM-SC isoforms respectively. α-HSP70 was used to indicate equal loading and for normalization. (F, G) Quantification of Western blots shown in (E) and expressed as relative expression (highest value = 1). Data are mean ± SEM of n=2 biological replicates. Technical duplicates were performed for each biological replicate for mRNA analysis. Asterisks denote significant differences between LP3 and SP5 or LP3 and SP10 at each ZT: ***p<0.001, **p<0.01, *p<0.05, two-way ANOVA with *post-hoc* Tukey’s HSD tests. (F) p(LP3)=1, p(SP5)<0.05, p(SP10)=0.005, JTK-CYCLE.

**Figure S5.**
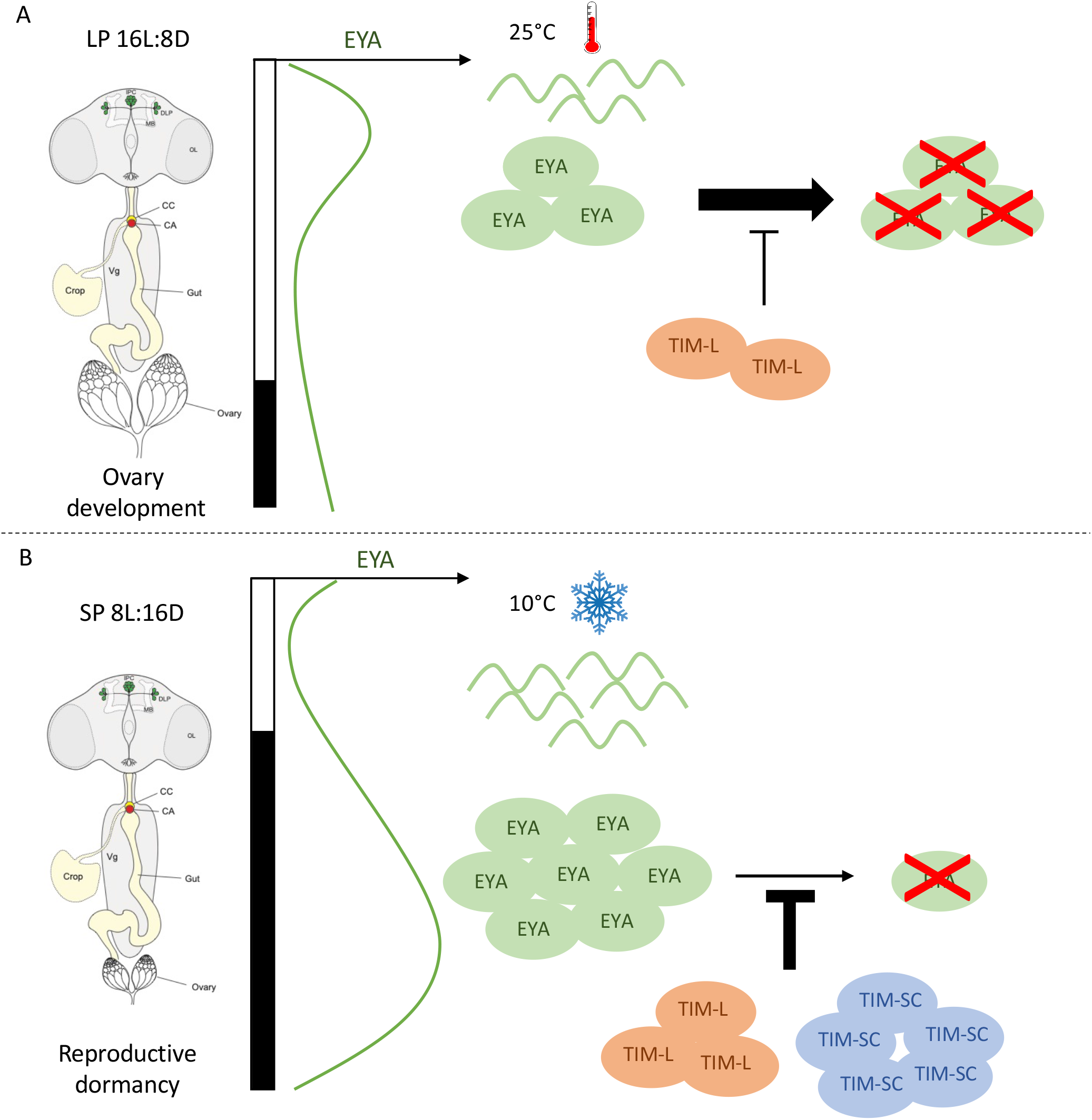
Schematic model of the roles of EYA and TIM on incidence of reproductive dormancy in response to temperature and photoperiodic cues. (A) In LP 16L:8D, 25°C (summer-like condition), flies are reproductively active and exhibit fully developed ovaries. In these conditions, *eya* mRNA expression is moderate but relatively low level of TIM-L is not sufficient to stabilize EYA at the scotophase at 25°C, restricting its moderate expression to the photophase. (B) In SP 8L:16D, 10°C (winter-like condition), *eya* mRNA is enhanced in the cold due to yet unknown mechanism. Concomitantly increase in TIM-L under SP coupled to TIM-SC induction at low temperature promotes EYA protein stabilization and accumulation in the scotophase, resulting in increased incidence of reproductive dormancy.

**Figure S6.**
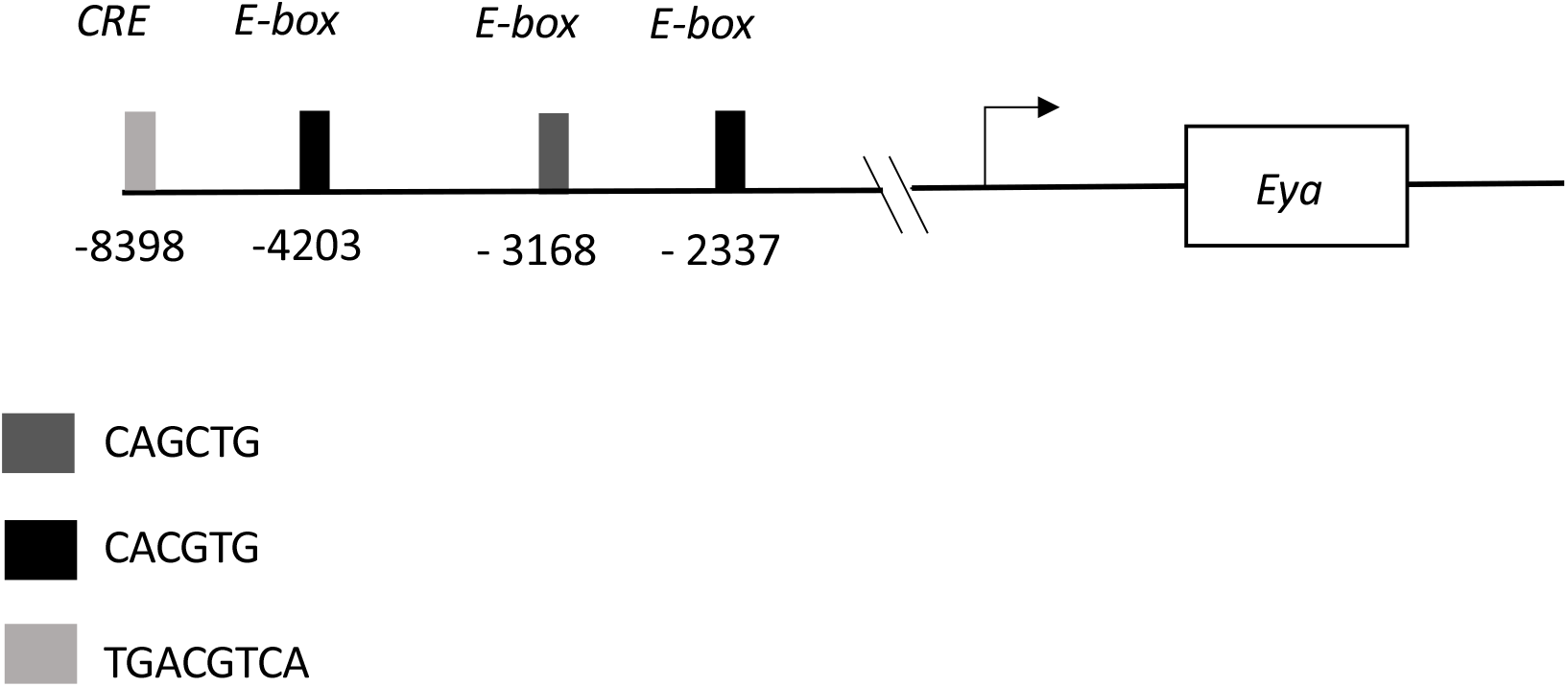
Upstream regulatory region of the *eya* gene. Analysis of 10kb upstream sequence from start codon, performed using MAFT. Black and Grey rectangles represent the position of E-boxes, light grey rectangle represent the position of CREB sequence from the start codon represented by a black arrow (3).

**Figure S7.**
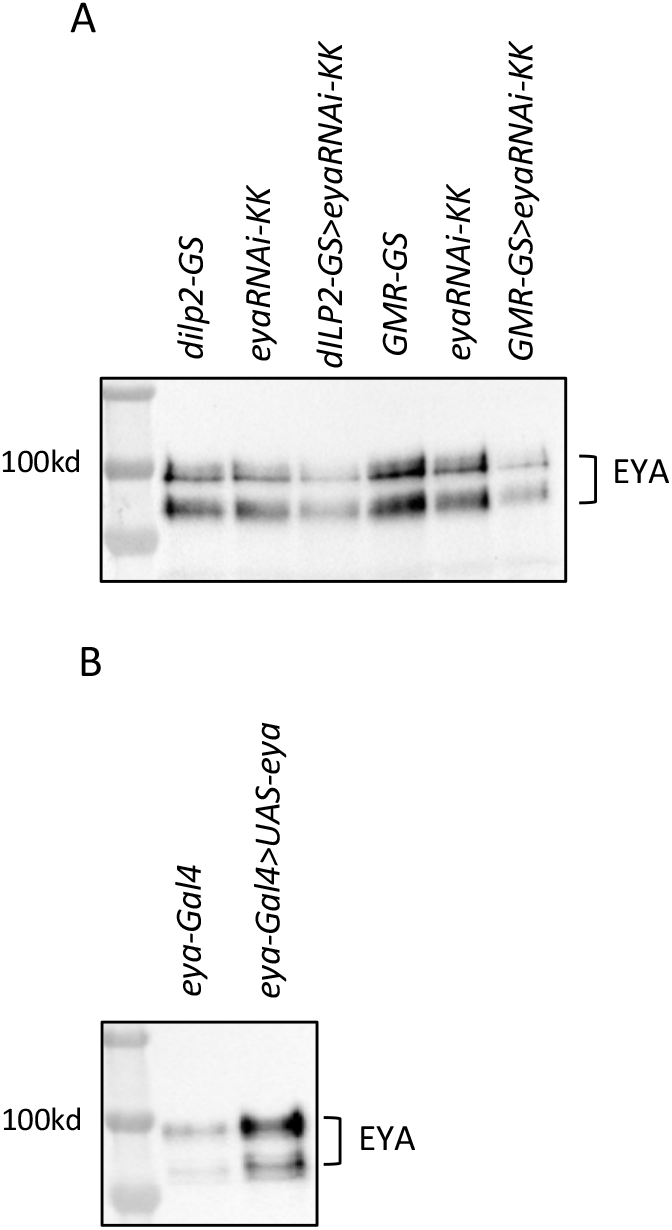
Validation of *eya* RNAi knockdown and overexpression in flies. (A) Western blot comparing EYA expression between parental control lines and flies expressing *eya* dsRNA in *dilp2* neurons using *dilp2-GS* gene-switch driver or in the visual system, using *gmr-GS* gene-switch drivers crossed with *UAS-eyaRNAiKK* responder line. (B) Western blot comparing EYA expression between parental control (*eya-Gal4*) and flies overexpressing EYA (*eya-Gal4>UAS-eya*).

## SI Materials and Methods

### Fly stocks

All stocks were in *w*^*1118*^ background except for *yw*, *tim*^*0*^, and were raised on Bloomington standard fly media at room temperature. Immunocytochemistry was performed using *w*; *eya-Gal4* (BDSC no.49292) or *w*; *dilp2-Gal4* (BDSC no.37516) crossed with *w*; *UAS-cd8-GFP* line (BDSC no. 32184). Genetic manipulation of *eya* expression was achieved using the UAS/Gal4 (4) and inducible gene-switch systems (5). To enhance *eya* endogenous expression levels, *w*; *eya-Gal4* (BDSC stock no. 49292) was crossed with *w*; *UAS-eya* line from the Bloomington *Drosophila* Stock Center (BDSC stock no. 5676). Targeted overexpression in *dilp2* expressing neurons was achieved using *w*; *dilp2-GS-Gal4* (kind gift from Henri Jasper) and in the eyes using *w*; *gmr-GS-Gal4* (BDSC no. 6759). Targeted expression of *eya*-RNAi in these cell types were similarly achieved. Two independent responder RNAi lines from the Vienna *Drosophila* Resource Center (VDRC stock no.107568 and 43911) were tested. Results were obtained using line no.107568 (*UAS-eya-RNAiKK* in the manuscript). The parental line no. 43911 showed reduced levels in *eya* expression in absence of induction and was therefore not included in the study. Induction of the gene-switch driver was performed as previously described in McGuire et al. (6) Newly emerged females used for gene expression analysis and diapausing assay were transferred to media supplemented with RU486 (final concentration 160ug/ml of food, dissolved in 100% EtOH) in order to induce the expression of *eya* or *eya* dsRNA after emergence and prevent the complication due to eye developmental defects. Control flies were transferred to media supplemented with EtOH to validate the specificity of the treatment. Validation of EYA overexpression and RNAi-induced knockdown at the protein level was performed by Western blotting (Fig. S7).

To assay the effect of *tim* on *eya* expression, we used *yw*; *tim*^*0*^ and *w*; *p{tim(WT)-3XFLAG}* and compared them to controls with appropriate genetic backgrounds. A *p{tim(WT)-luc}* transgene, containing 4.5kb of *tim* promoter, *tim*-*ls* full-length coding region, and a luciferase reporter in *pattB* vector, was kindly provided by Patrick Emery. The luciferase reporter was removed using XhoI/MluI restriction sites and 3XFLAG-6XHIS epitope was added in frame to the N-terminus of the TIM coding region. Plasmids were injected into *w*^*1118*^ fly embryos carrying *attP* sites on chromosome 3 (*attP2*) (BestGene, Inc., Chino, CA) and overexpression of TIM was evaluated by Western blotting.

### Reproductive dormancy assay

Newly emerged females were reared in short photoperiod (SP 8L:16D) or long photoperiod (LP 16L:8D) at 10°C for 28 days in DigiTherm temperature-controlled incubators (Tritech Research Inc., Los Angeles, CA) prior to ovary dissection. LED was used as source of light to minimize variation in temperature between SP and LP due to difference in day length. Temperatures were recorded using HOBO^®^ meter (Onset Computer Corporation, Bourne, MA) and no more than +/−0.4°C variation in average between light and dark phases were observed. Reproductive dormancy in WT and *eya* overexpressing flies were also evaluated in temperature adjusted incubators in which the temperature was set at 9.2°C during the light phase to abolish variations in temperature between light and dark phases (Fig. S1). Levels of reproductive dormancy (n=30-40 individuals per group) was determined by measuring ovary surface size (expressed in pixel^2^) using the EVOS microscope (Life technologies, Carlsbad, CA) in combination with image J software.

### Photoperiodic and temperature shift time-course experiments

To determine the role of photoperiodic changes in *eya* expression, newly emerged flies were reared at 10°C for 3 days in long photoperiod, 16 hrs Light:8 hrs Dark (LP 16L:8D) and then for 7 days in short photoperiod, 8 hrs Light:16 hrs Dark (SP 8L:16D). Flies were collected every 4h over a 24h period on days 3, 5 and 10 labeled LP3, SP5 and SP10 respectively (Fig. 2A). For temperature shift experiments, flies were reared in 12 hrs light:12 hrs dark LD cycles (LD 12:12) at 25°C for 3 days (3D25). After day 3, one group of flies remained at 25°C (10D25) while the second group was transferred to 10°C (10D10) for 7 additional days. Samples were collected at day 3 (prior to temperature shift) and day 10 (Fig. 3A). All samples were collected and flash frozen on dry ice and stored at −80°C prior to RNA or protein extraction.

### Whole-Mount Brain Immunocytochemistry

To assess overall *eya* expression pattern in the brain (Fig. S2), three-day old adult flies were entrained to a 12:12 LD cycle at 25°C for 4 days and dissected on the 5th day at indicated time-points. Flies were first fixed in 4% formaldehyde for 45-60 min at room temperature and were subsequently dissected in 1X Phosphate-Buffered Saline with 0.1% Triton-X (1XPBST). Brains were quickly washed three times (1XPBST) and followed by 3 × 10 min (1XPBST) washes on a tube rotator, then blocked for 90 min in 10% Normal Goat Serum in PBST at room temperature. Brains were incubated in primary antibody (Mouse α-GFP, 1:200, Invitrogen MA5-15349) overnight at 4°C and rinsed 6 × 20 min in 1XPBST. Next, they were incubated with secondary antibody (α-mouse-AF488, 1:200, Jackson immunoresearch, West Grove, PA) overnight at 4°C, rinsed 6 × 20 min in 1X PBST, and mounted with Vectashield (Vector Laboratories, Burlingame, CA). Brains were imaged using a Leica TCS SP8 confocal microscope. For EYA and GFP double staining to determine colocalization in IPCs (Fig. 1B), flies were collected on LP3 and SP5 (ZT16) at 10°C similar to photoperiodic shift experiment. Mouse α-EYA10H6 (deposited to the DSHB by S. Benzer and N. M. Bonini) was diluted at 1:5 and chicken α-GFP (Invitrogen, A10262) was used at 1:200 dilution. Secondary antibodies were α-mouse-Cy3 and α-chicken-AF488 (both used at 1:200, Jackson immunoresearch). EYA levels were quantified using image J. For TIM and GFP double staining (Fig. 7C-E), flies were collected on SP5 (ZT16) at 10°C. Rat α-TIM (N-terminal antigen, M. Young) was diluted at 1:300 and chicken α-GFP (Invitrogen, A10262) was used at 1:200 dilution. Secondary antibodies were α-rat-Cy3 and α-chicken-AF488 (both used at 1:200 (Jackson immunoresearch).

### Analysis of circadian/daily gene expression by real-time PCR

Flies were entrained for 3 days in LD (12:12) and collected every 4 h on the 4th day. cDNA was generated using Superscript IV (Life Technologies) and real-time PCR was performed using SsoAdvanced SYBR green supermix (Bio-Rad) in a CFX96 (Bio-Rad). Primers were designed to detect these transcripts: for *eya:* 5’-GAGGCCTGGCTACAGATACG-3’ and 5’-AGTTGCGTGGAGGTTACCAG-3’; for *tim-L*: 5’-CCCTTATACCCGAGGTGGAT-3’ and 5’-TGATCGAGTTGCAGTGCTTC-3’; *timL+M*: 5’-CTCCATGAAGTCCTCGTTC-3’ and 5’-TGTCGCTGTTTAATTCCTTC-3’; *tim-M*: 5’-GGAGACAATGTACGGACTC-3’ and 5’-ATTTCACACAGAGAGAGAGC-3’, *tim-cold*: 5’-GCATCTGTGTACGAAAAGGA-3’ and 5’-ATGTAACCTATGTGCGACTC-3’; *tim-sc* 5’-AACACAACCAGGAGCATAC-3’ and 5’-ATGGTCCACAAATGTTAAAA-3’. *cbp20* (5’-GTCTGATTCGTGTGGACTGG-3’ and 5’-CAACAGTTTGCCATAACCCC-3’) was used for normalization. Primers for amplification of *tim* splice variants were described in (7). Cycling conditions were 95°C for 30 seconds, 40 cycles of 95°C for 5 seconds, followed by an annealing/extension phase at 60°C for 30 seconds. The reaction was concluded with a melt curve analysis going from 65°C to 95°C in 0.5°C increments at 5 seconds per step. Two technical replicates were performed for each biological replicate. Two to three biological replicates were performed for analysis of each gene as indicated. Data were analyzed using the standard ΔΔCt method and target gene mRNA expression levels were normalized to the reference gene mRNA levels (*cbp20*), which remain unchanged over a day regardless of temperature and photoperiodic changes. Finally, Ct values for all time-points were divided by the highest Ct value to generate a scale from 0 to 1 to indicate relative expression.

### Western blotting and antibodies

5% blocking reagent (Bio-Rad, Hercules, CA) in 1XTBST was used for incubation with all antibodies. Primary antibodies for western blotting analysis were used at the following dilution: mouse α-EYA10H6 (DSHB) at 1:1000, rat α-TIM (Chiu lab, RRID:AB_2782953, C-terminal antigen aa1133-1421) at 1:1000, rat α-TIM (kind gift form Michael Young and Deniz Top, N-terminal antigen aa222-557 (1, 2) at 1:1000, mouse α-HSP70 (Sigma, St. Louis, MO) at 1:7000. The levels of HSP70 (loading control) were found to be stable between 10°C and 25°C (see Fig. 6A). Secondary antibodies were used as follow: α-mouse-IgG-HRP (GE Healthcare, Piscataway, NJ) at 1:1000 for α-EYA10H6 and 1:7000 for α-HSP70 detection, α-rat-IgG-HRP (GE Healthcare) at 1:1000 for both α-TIM. Membranes were imaged and protein levels were quantified using the ChemiDoc MP system with Image Lab software (Bio-Rad, Hercules, CA).

### Coimmunoprecipitation (CoIP) in Drosophila S2 cells

3 × 10^6^ *Drosophila* S2 cells were transiently transfected with pAc-*eya*-3XFLAG-6XHis in combination with either pAc-*tim-l*-3HA or pAc-*tim-sc-*3HA and reciprocal coIP assays were performed as described in (8).

### Statistics

Cycling statistics (rhythmicity, period, amplitude and peak phase) of daily mRNA and protein expression were determined using the program JTK-CYCLE in R (9). Other statistical analyses were performed using GraphPad Prism 8.0 (GraphPad Software, La Jolla California USA). In the case of normally distributed data (Shapiro-Wilk normality test, p>0.05), statistical significance of differences in daily mRNA and protein abundance was tested by two-way ANOVA with *post-hoc* Tukey’s HSD tests. If data was not normally distributed, non-parametric Mann Whitney was applied. The mean of technical replicates was used to analyze differences between biological replicates. Error bars = standard error of the mean (SEM) of biological replicates. Asterisks indicate significant differences in abundance between genotypes or conditions at indicated time-points. Differences in ovary size were analyzed using one- or two-way ANOVA followed by *post hoc* Tukey’s multiple comparisons tests. If data was not normally distributed, non-parametric Kruskall-Wallis test with Dunn’s multiple comparison was applied. Error bars = SEM. For coimmunoprecipitation assays, variances in protein interactions were analyzed using two tailed Student’s test. Error bars = SEM.

### PCR for timeless genotyping

A pool of ten individuals per fly strain was used to extract genomic DNA as described in (10). The *tim* region containing the polymorphic site was amplified and sequenced using the following forward and reverse primers *tim*(F): 5’-GTGGTTGCGTAATGCCCTGGAAGA-3’ and *tim*(R): 5’-TCACCTCCTGCAAAGTGGCC-3’.

---

**Table S1.** The *timeless* background of the *Drosophila* strains used in reproductive dormancy assays, determined by sequencing. *ls* = long and short allelic variant; s = short allelic variant (11, 12).

| *Genotype* | *timeless variant* |
| --- | --- |
| *w*<sup>1118</sup> | *s/s* |
| *w; dilp2-GS-Gal4* | *s/s* |
| *w; gmr-GS-Gal4* | *ls/ls* |
| *w; UAS-eyaRNAikk* | *ls/ls* |
| *w; dilp2-GS-Gal4>w; UAS-eyaRNAikk* | *s/ls* |
| *w; gmr-GS-Gal4>w; UAS-eyaRNAikk* | *ls/ls* |
| *w; eya-Gal4* | *s/s* |
| *w; UAS-eya* | *s/s* |
| *w; eya-Gal4>w; UAS-eya* | *s/s* |
| *w; dilp2-GS-Gal4>w; UAS-eya* | *s/s* |
| *w; gmr-GS-Gal4>w; UAS-eya* | *ls/s* |
| *yw* | *s/s* |
| *w; p{tim(WT)-3XFLAG}* | *ls/ls* |
| *yw; tim*<sup>0</sup> | X |

